# Molecular Profiling and High-Content Drug Screening of Metastatic Colorectal Cancer Organoids Reveal Evolutionary Mechanisms of Pan-KRAS Inhibitor Resistance

**DOI:** 10.64898/2026.01.14.699345

**Authors:** Gizem Damla Yalcin, Kubra Celikbas Yilmaz, Raffaella Klima, Olcay Kurtulan, Duriye Ozer Turkay, Oktay Halit Aktepe, Ahmet Bulent Dogrul, Cenk Sokmensuer, Volkan Oter, Erdal Birol Bostanci, Javier Fernandez-Mateos, Suayib Yalcin, Luca Braga, Ahmet Acar

**Affiliations:** Department of Biological Sciences, Middle East Technical University, Universiteler Mah. Dumlupınar Bulvarı No: 1, 06800 Çankaya, Ankara, Türkiye; Functional Cell Biology Group, International Centre for Genetic Engineering and Biotechnology (ICGEB), 34149 Trieste, Italy; Department of Pathology, Faculty of Medicine, Hacettepe University, Sıhhiye, 06100 Ankara, Türkiye; Department of Pathology, Ankara City Hospital, Ankara, Türkiye; Department of Medical Oncology, Dokuz Eylul University, Izmir, Türkiye; Department of General Surgery, Faculty of Medicine, Hacettepe University, Sıhhiye, 06100 Ankara, Türkiye; Department of Gastrointestinal Surgery, Ankara City Hospital, Ankara, Türkiye; Department of Biochemistry and Molecular Biology, University Insitute of Oncology of Asturias, University of Oviedo, Oviedo, Spain; Department of Medical Oncology, Faculty of Medicine, Hacettepe University, Sıhhiye, 06100 Ankara, Türkiye

## Abstract

Metastatic colorectal cancer exhibits extensive intertumoral and intratumoral heterogeneity, which underlies variable treatment responses and the inevitable emergence of drug resistance. Preclinical models that accurately reflect this heterogeneity and enable the investigation of resistance mechanisms at the clonal level, while also providing sensitive and quantitative measurements of treatment response, remain limited. Here, we describe a living biobank of patient-derived organoids (PDOs), generated from metastatic colorectal cancer samples encompassing various genetic backgrounds, including different KRAS genotypes. These PDOs retain key phenotypic, genomic, and transcriptional features of their matched tumors, and, when coupled with a multiparametric high-content 3D drug-screening platform integrating volumetric growth, nuclear content, and proliferative readouts, enable sensitive detection of subtle responses that are not captured by conventional viability-based assays. Systematic phenotypic profiling identified KRAS mutational status as a major determinant of drug response, with KRAS wild-type organoids exhibiting predominant cytotoxic collapse, whereas KRAS-mutant PDOs displayed cytostatic growth arrest. Building on this phenotypic stratification, we investigated resistance to the pan-KRAS inhibitor BI-2865 by integrating longitudinal drug exposure with expressible DNA barcoding and single-cell RNA sequencing to resolve clonal and transcriptional dynamics. Resistance to BI-2865 arose primarily through transcriptional plasticity rather than widespread genomic alterations. Lineage tracing revealed genotype-dependent trajectories, with de novo emergent clones driving resistance in KRAS^WT^ PDO, whereas KRAS^G13D^ models were dominated by expansion of pre-existing clones. Single-cell analysis on KRAS^G13D^ demonstrated a shared transcriptional landscape comprising multiple resistant states rather than a single convergent identity. Together, our findings define resistance to pan-KRAS inhibition as a modular, state-resolved process driven by transcriptional plasticity and genotype-dependent clonal dynamics and establish PDO-based multiparametric phenotyping combined with lineage-resolved single-cell profiling as a robust framework for dissecting the evolutionary framework of therapeutic resistance in metastatic colorectal cancer.

## Introduction

As one of the most frequently diagnosed cancers, colorectal cancer (CRC) accounts for a substantial proportion of cancer deaths worldwide, largely due to the high incidence of metastasis in more than half of the patients^1^. While chemotherapy and targeted therapy agents have led to improved clinical outcomes, treatment resistance is inevitable particularly due the prevalence of both inter- and intra-tumoral heterogeneity which continue to hinder durable therapeutic success^2,3^. To effectively investigate tumor heterogeneity and therapy resistance, there is a critical need for physiologically relevant preclinical models, such as patient-derived organoids (PDOs), that enable the identification of predictive biomarkers and therapeutic vulnerabilities.

Over the past decade, PDOs have gained considerable attention as successful preclinical models for studying therapeutic response. In contrast to conventional two-dimensional cell lines, PDOs retain key genetic, phenotypic and functional characteristics of their source tumors, serving as a suitable platform for studying tumor heterogeneity and treatment response^4–7^. In CRC^8^ and across different solid malignancies^9–12^, PDOs have been demonstrated to recapitulate specific genomic alterations and drug response patterns, highlighting their potential use as a predictive model in personalized medicine^13,14^. With such features, PDOs enabled the establishment of living biobanks and designing of co-clinical trials in a scalable ex vivo platforms to assess the systemic evaluations of therapeutic vulnerabilities across genetically diverse patient cohorts. These advancements positioned PDOs as a critical pre-clinical model for both translational research and personalized medicine.

Despite KRAS mutations being a well-known oncogenic driver in the progression of CRC, the molecular mechanisms underlying resistance to KRAS-targeted therapies, particularly next-generation pan-KRAS inhibitors, remain insufficiently characterized^15,16^. Growing evidence supports the concept that therapeutic resistance frequently emerges through Darwinian selection^17–19^. In this process, pre-existing or drug-induced subclonal populations with a survival advantage under treatment pressure undergo positive selection^20^. This clonal evolutionary process reflects the inherent plasticity and intratumoral heterogeneity of malignant cell populations and highlights the adaptive fitness landscapes that tumors can exploit to evade targeted interventions^21^. Elucidating these evolutionary dynamics, especially in a model system effectively recapitulating the dynamics of patients’ tumors, is therefore critical for anticipating resistance trajectories and identifying rational combination strategies that can preempt or overcome therapeutic failure.

In this study, we generated a living PDO biobank from 20 metastatic CRC patients and performed extensive molecular and phenotypic characterizations including H&E staining, immunohistochemistry (IHC), whole-exome sequencing (WES), and bulk RNA sequencing. To functionally interrogate therapeutic vulnerabilities, we established a high-content, 3D image-based drug screening platform using three distinct fluorescent markers marking total organoid volume, nuclei number and EdU+ statues under 50 number for small molecule inhibitors. This enabled spatially resolved, multiparametric profiling of drug responses, capturing inter-patient drug response at high resolution. To investigate mechanisms of acquired drug resistance, we selected two genetically defined PDO models, harboring KRAS^WT^ and KRAS^G13D^ genotypes, from our biobank and exposed them to the novel pan-KRAS inhibitor BI-2865, a selective, non-covalent small-molecule inhibitor that binds to the inactive GDP-bound form of KRAS thereby preventing its activation and downstream signaling^22^. Given the limited current understanding of resistance mechanisms to this compound, these models offered a controlled experimental platform to dissect molecular and phenotypic adaptations associated with therapeutic escape in both KRAS-mutant and wild-type settings. We performed integrative molecular profiling of pre- and post-resistant PDOs using single-cell RNA sequencing, WES, bulk RNA-seq, and 3D image-based drug screening. In parallel, we implemented a high-complexity cellular barcoding strategy to track clonal dynamics of resistance acquisition in these PDOs, enabling reconstruction of evolutionary trajectories and identification of resistant subpopulations selected under drug pressure. Together, these integrative approaches present both inter-patient variability in therapeutic response and the evolutionary principles governing KRAS inhibitor resistance, providing a framework for the development of more effective precision treatment strategies in metastatic CRC.

## Results

### Establishment and molecular validation of metastatic CRC PDOs

We established a PDO biobank from surgically resected liver metastases obtained from 19 patients with CRC and 1 gastric cancer, serving a representative in vitro resource of metastatic disease. Following tumor resection, metastatic tissues were subjected to combined enzymatic and mechanical dissociation, and viable epithelial fractions were expanded under 3D culture conditions. To rigorously assess the biological resemblance of each PDOs to their source tumor tissue samples, we performed integrative multi-layer profiling on all 20 matched PDO–tumor pairs. This comprehensive characterization included histomorphological assessment, WES, bulk RNA sequencing, and high-content 3D image-based drug response profiling **(Fig. 1a)**, enabling simultaneous evaluation of phenotypic, molecular and functional heterogeneity. The concordance between PDOs and their tumors of origin was evaluated using H&E staining and assessed by two independent pathologists, conforming preservation of key architectural and morphological features across all matched pairs **(Fig. 1b)**. Immunohistochemical analyses were performed across all 20 PDOs. Ki-67 staining revealed variable proliferative activity among PDOs, consistent with the presence of PDOs exhibiting distinct growth rates. In contrast, expression of the intestinal lineage marker CDX2 and the epithelial marker CK20 was consistently preserved across the PDO models **(Fig. 1b).**

**Figure 1.**
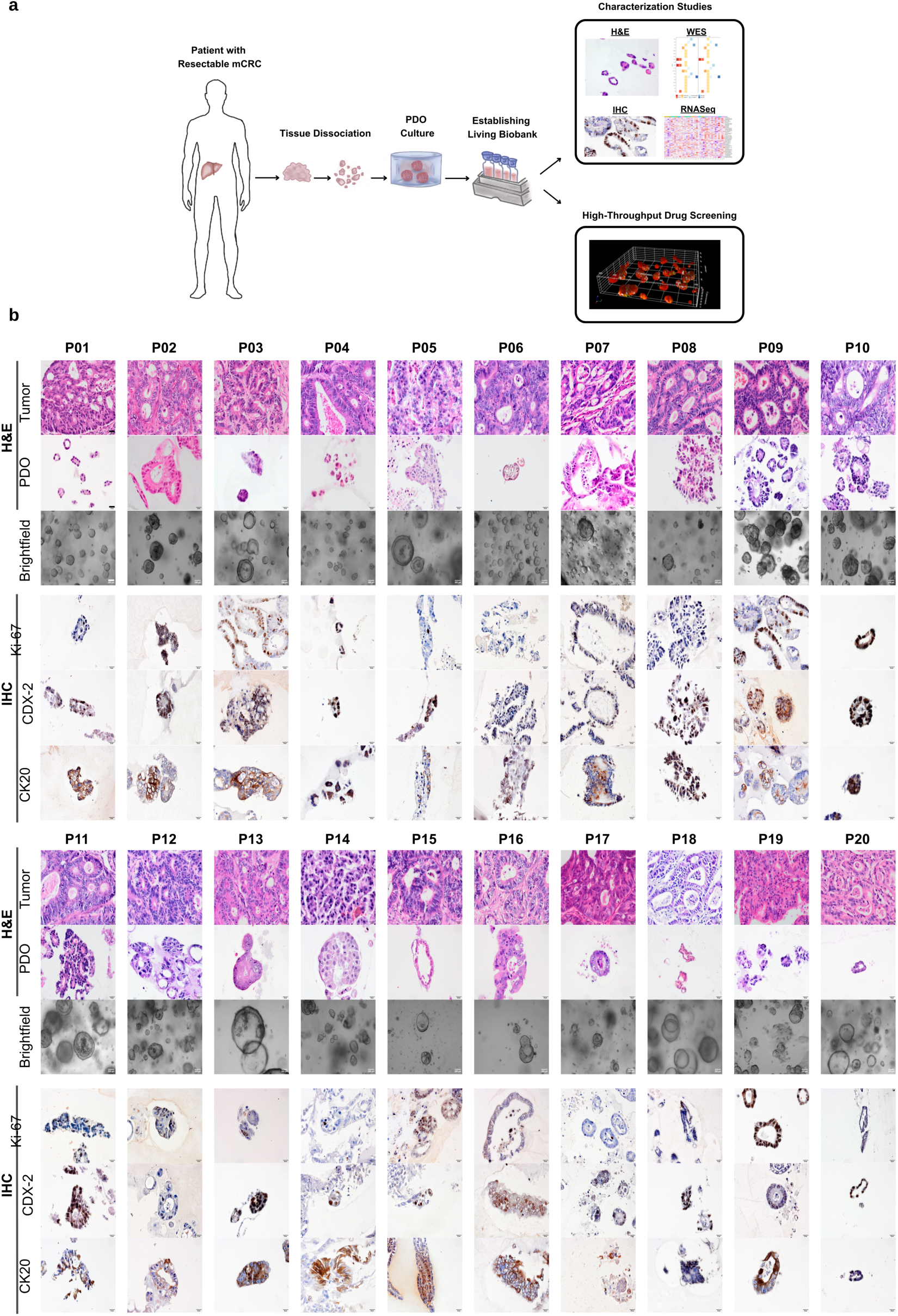
Establishment and histological validation of metastatic colorectal cancer organoids. **a** Schematic overview of PDO generation from surgically resected mCRC. Tumor specimens were enzymatically and mechanically dissociated and expanded under 3D culture conditions to establish a patient-matched living PDO biobank. PDOs were characterized by H&E staining, IHC, WES, and bulk RNA-seq, followed by high-throughput 3D image–based drug screening**. b** Histological comparison of matched tumors and representative PDOs from 20 patients (P01–P20). H&E-stained parental tumors (top) display colorectal adenocarcinoma morphology, while corresponding PDOs (middle) retain glandular architecture and epithelial polarity. Brightfield images (bottom) highlight inter-patient variability in PDO growth. Tumor and PDO H&E images were acquired at 40× (scale bar, 20 µm; Olympus CellSens), and brightfield images at 10× (scale bar, 200 µm; AMScope). IHC analysis of PDO sections from patient PDO-01-PDO-20 showing Ki-67, CDX2, and CK20 staining, confirming preservation of proliferative capacity and intestinal epithelial identity. Scale bar, 50 µm (20×; Olympus CellSens).

Next, we interrogated whether PDOs retained the genomic landscape of their parental tumors by performing WES on all 20 matched PDO–tumor pairs. Detailed SNVs for each tumor and corresponding PDOs are provided **(Supplementary Table 1).** Clinical metadata, including patient sex, primary tumor location, and prior treatment history, were integrated with genomic landscape **(Fig. 2a)**. Comparative mutational analysis of key CRC driver genes, including *APC*, *TP53*, *KRAS*, *PIK3CA*, and *SMAD4*, revealed a high degree of concordance in somatic mutation profiles between tumors and their corresponding PDOs, with the overall molecular landscape closely resembling that reported for metastatic CRC cancers in the previous studies^23,24^.

**Figure 2.**
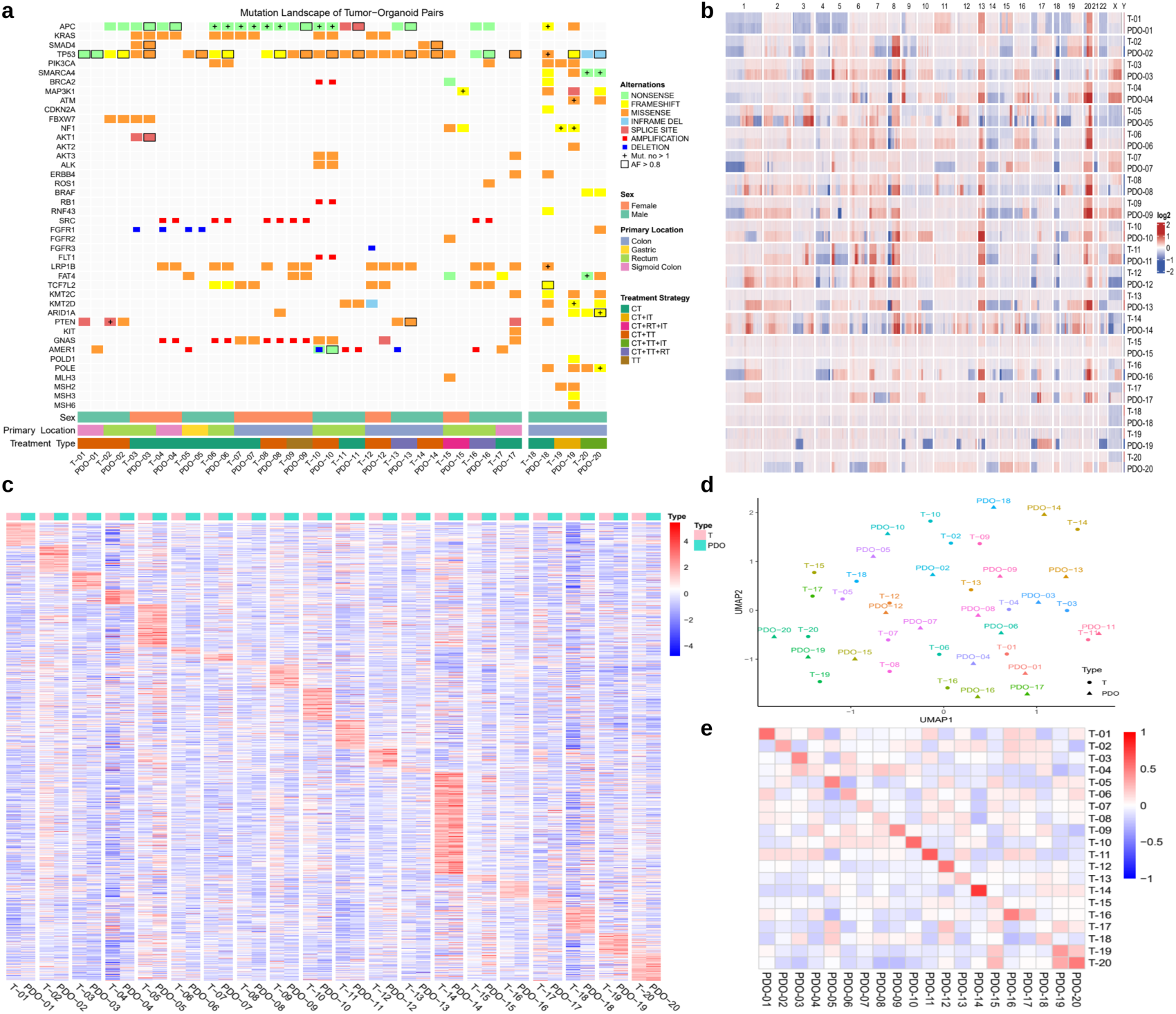
Genomic and transcriptomic characterization of matched tumor–PDO pairs. **a** Mutation landscape of 20 matched tumor–PDO pairs. The oncoplot summarizes nonsynonymous alterations in recurrent colorectal cancer genes. Variants are classified by type (nonsense, frameshift, missense, in-frame deletion, splice-site, copy-number gain or loss). High-allele-frequency variants (AF > 0.8) are indicated by square outlines, and loci with multiple mutations are marked by plus signs. Clinical annotations include sex, primary tumor location, and prior treatment strategy (CT, chemotherapy; IT, immunotherapy; TT, targeted therapy; RT, radiotherapy; combined regimens indicate multi-modality treatment). WES was performed for each tumor–PDO pair, and high-confidence somatic variants were retained after filtering against matched blood DNA. **b** Copy-number variation profiles across 20 matched tumor–PDO pairs. Heatmap shows log2 copy-number ratios derived from WES across chromosomes 1–22 and X/Y for tumors (T) and corresponding PDOs. Red and blue denote relative gains and losses after normalization to matched blood DNA. **c** Bulk RNA-seq heatmap of 20 tumor–PDO pairs. Variance-stabilized log2 expression values for the 8,000 most variable genes are shown, with rows representing genes and columns representing samples. Values are centered and scaled per gene; red and blue indicate relative up- and downregulation. **d** UMAP of bulk RNA-seq profiles from matched tumor–PDO pairs. Each point represents a tumor or its corresponding PDO, colored by patient identity. Tumor–PDO pairs cluster in close proximity, indicating preservation of major transcriptomic features in organoid culture. **e** Pairwise transcriptomic correlation matrix of tumor–PDO pairs. Heatmap shows Pearson correlation coefficients computed from variance-stabilized log2 expression values across the 8,000 most variable genes, with stronger positive correlations indicated in red.

Notably, the mutational landscape across all the PDOs displayed inter-patient variability, consistent with the molecular heterogeneity that characterizes metastatic CRC **(Fig. 2a)**. Despite the overall concordance, three MSI cases (P-18, P-19, and P-20) showed partial overlap between tumor and PDO mutational profiles. MSI status in all three patients was confirmed from the primary tumor, yet the PDOs displayed additional mutations not detected in the tumor samples. Similar findings in MSI CRC organoids have been linked to the acquisition of new variants due to MMR deficiency ^25^ and to the selective expansion of minor subclones present in spatially heterogeneous tumors^26^. These findings demonstrate the successful establishment of PDOs capturing patient-specific histomorphological features and single-nucleotide variations while preserving inter-patient diversity. Chromosome-level CNV profiling demonstrated strong concordance between tumors and their matched PDOs across the cohort, with each pair exhibiting highly similar genome-wide patterns of copy number gains and losses. Notably, these alterations were consistent with CNV landscapes reported in primary colorectal cancer and liver metastatic CRC studies^27,28^, such as gains on 20q, 7, 8q, and 13q and losses on 18, and 4q **(Fig. 2b)**. Moreover, copy number gains in *SRC* in PDO-04, PDO-06, PDO-08, PDO-09, and PDO-16, and in *GNAS* in PDO-04, PDO-06, PDO-08, and PDO-09 were detected. *BRCA2*, *RB1*, and *FLT1* gains were confined to PDO-10, whereas *AMER1* amplification occurred only in PDO-11. A consistent copy number loss was identified for *FGFR1* in PDO-05, and all of these alterations were concordant with the profiles observed in the respective parental tumors **(Fig. 2a).** Detailed CNV profiles for each tumor–PDO pair are provided **(Supplementary Fig. 1).** Notably, extensive copy number alterations in MSS cases, reflected by broad chromosomal gains and losses across multiple loci, whereas MSI samples displayed minimal or no CNV changes, consistent with the known predominance of point mutations and indel events rather than arm-level copy number instability in MMR-deficient tumors^29,30^. Detailed patient characteristics, including MSI status for all patients, are provided **(Supplementary Table 2)**. This pattern confirms that PDOs faithfully recapitulate the chromosomal structural landscape of their parental tumors, while also highlighting the distinct genomic architecture of MSS-versus MSI-driven metastatic CRC **(Fig. 2b)**. We then asked the question whether retaining of morphological and genomic features are extended to transcriptional programs. To address this, we performed bulk RNA sequencing analysis on all matched tumor–PDO pairs. After filtering lowly expressed genes and selecting the 8.000 most variable transcripts, results revealed that PDOs closely recapitulate the global gene expression profiles of their tumor counterparts **(Fig. 2c)**. Despite clear inter-patient variability, each tumor-PDO pair consistently clustered together, indicating patient-specific transcriptional identity **(Fig. 2c)**. Dimensionality-reduction analyses using UMAP further supported this finding, with matched tumors and PDOs positioning in close proximity despite separation across patients **(Fig. 2d)**. Similarly, correlation analysis demonstrated that each tumor clustered most closely with its corresponding PDO, indicating high transcriptomic fidelity across all of the PDOs **(Fig. 2e)**. Together, these integrated histological, genomic and transcriptomic analyses demonstrate that PDOs faithfully preserve both the molecular identity and inter-patient heterogeneity of metastatic colorectal cancer. These findings establish a robust and clinically relevant foundation for downstream functional drug screening and mechanistic interrogation of therapeutic response and resistance.

### Differential drug response profiles in KRAS-Wild-Type and KRAS-Mutant PDOs

Having established that the PDOs successfully recapitulates the genetic and transcriptional landscape of the original tumors, we next leveraged this platform to functionally interrogate therapeutic vulnerabilities using high-throughput drug screening approach. Functional drug response profiles were generated for 10 PDOs using a high-content, 3D imaging platform. Drug sensitivity was quantified across three complementary phenotypic parameters: organoid volume (V), total nuclei count (N), and the proportion of EdU-positive nuclei (E), reflecting proliferative activity. Image acquisition and stepwise 3D segmentation, feature extraction, and quantitative analysis were performed using the Harmony high-content analysis software. A curated panel of 50 compounds, tested at a single dose of 1 µM, targeting diverse oncogenic and cellular pathways, including cell-cycle regulation, PI3K/AKT/mTOR, MAPK, DNA damage response, tyrosine kinases, stem cell and WNT signaling, and TGF-β pathway, was screened across the PDO cohort **(Fig. 3a)**. To enable objective and biologically meaningful hit identification, we implemented a two-tiered, phenotype-integrated classification strategy. Compounds were classified as cytotoxic when they induced a concurrent reduction in organoid volume (V), nuclei count (N), and EdU incorporation (E), or when the combined decrease in volume and nuclei exceeded 50%, reflecting a transition from growth inhibition to structural disintegration and cell death. In contrast, cytostatic responses were defined by preserved structural integrity (V, N > 50%) accompanied by a marked suppression of proliferative activity (E < 50%), indicative of cell-cycle arrest without overt cell death. This multi-parametric, single-organoid-resolved hit-calling strategy, based on object-level measurements, enables precise discrimination between cytostatic and cytotoxic drug effects. Collectively, this imaging-based screening platform provides a highly quantitative and biologically informative framework to resolve subtle yet therapeutically relevant drug sensitivities that are not captured by conventional bulk viability assays.

**Figure 3.**
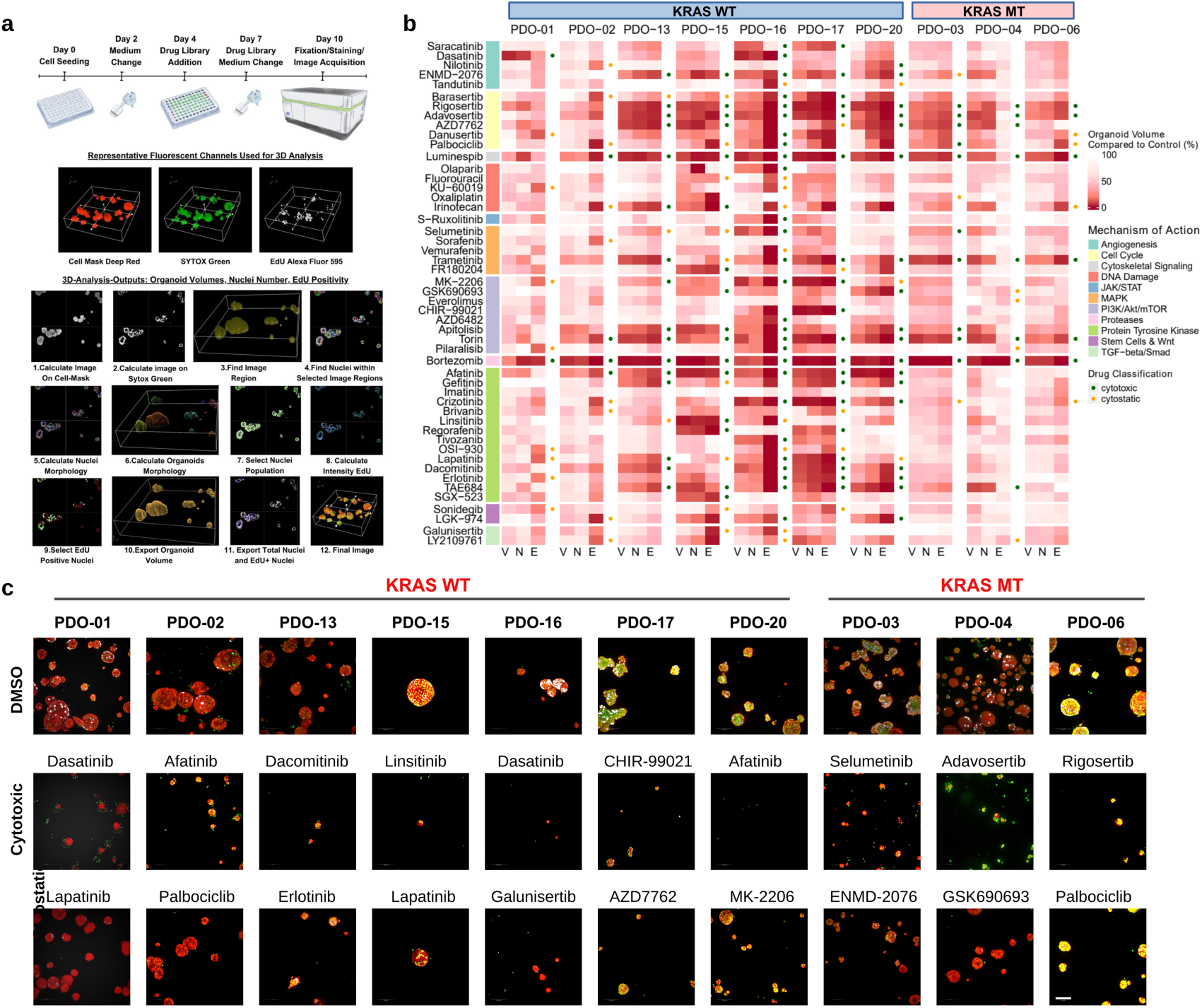
Multiparametric high-throughput drug response profiling in patient-derived organoids. **a** Schematic of the 10-day high-throughput 3D drug screening workflow. PDOs were seeded in 384-well plates (day 0), allowed to form organoids for 4 days, and treated with a targeted compound library from day 4 to day 10, with re-dosing on day 7. On day 10, organoids were fixed and stained with SYTOX™ Green Nucleic Acid StainS7020 (Life technologies)(488 nm HCS cell mask deep red, H32721 (Life technologies) 647 nm), and Click-iT EdU Cell Proliferation Kit for Alexa Fluor 594 C10339 (Life Technologies)594 to assess nuclear content, organoid volume, and proliferative activity. **b** Heatmap of multiparametric drug response profiles across 10 PDOs. Drug sensitivity was quantified using three phenotypic metrics: organoid volume (V), total nuclei count (N), and EdU-positive nuclei fraction (E). A panel of 50 compounds (1 µM, single dose) targeting diverse signaling pathways was screened, and responses are shown as reductions relative to DMSO controls. Compounds were classified as cytotoxic (green dots) when V, N, and E were jointly reduced or when combined loss of V and N exceeded 50%, and as cytostatic (yellow dots) when structural integrity was preserved (V and N > 50%) despite strong suppression of proliferation (E < 50%). **c** Representative 3D images of PDO responses under DMSO control, cytotoxic, and cytostatic conditions. Images show maximum projections from one of five fields per well, illustrating either structural collapse and loss of nuclei or preserved architecture with reduced EdU incorporation. Confocal Z-stacks were acquired on an Operetta CLS system (20× water-immersion objective, NA = 1.0; 1 µm Z-steps). Scale bar, 100 µm.

To resolve the impact of KRAS mutational status on therapeutic sensitivity, we next performed a KRAS-stratified analysis of cytotoxic and cytostatic drug responses. Across the ten metastatic CRC PDOs, stratified by KRAS status (KRAS-mutant: PDO-03, PDO-04, PDO-06; KRAS-wild-type: PDO-01, PDO-02, PDO-13, PDO-15, PDO-16, PDO-17, PDO-20), the single-dose (1 µM) 3D imaging screen identified a total of 204 phenotypic hits, comprising 144 cytotoxic and 60 cytostatic responses. Marked inter-patient variability was observed in the total number of hits per PDO, ranging 10 to 36, highlighting the substantial functional heterogeneity within the cohort **(Fig. 3b)**. This variability closely aligns with well-documented KRAS-driven differences in drug susceptibility in organoid models^31^ . Indeed, KRAS-mutant PDOs (PDO-03, PDO-04, and PDO-06) more frequently exhibited cytostatic or non-responsive profiles rather than cytotoxic responses to inhibitors (afatinib, dacomitinib, erlotinib, gefitinib, lapatinib) targeting the EGFR signaling pathway. This pattern mirrors prior organoid-based studies demonstrating that RAS-mutant CRC models often undergo proliferation arrest or display primary resistance, rather than apoptotic cell death, following EGFR blockade^32^.

Within the KRAS wild-type subgroup (PDO-01, PDO-02, PDO-13, PDO-15, PDO-16, PDO-17, and PDO-20), the majority of cytotoxic responses clustered around mechanisms targeting proteostasis, mitotic checkpoint control, receptor tyrosine kinase targeting and PI3K/mTOR signaling **(Fig. 3b)**. The proteasome inhibitor bortezomib elicited strong cytotoxic effects across all seven KRAS WT PDOs, indicating a broad dependency on intact protein homeostasis as previously reported^7,33^ . Similarly, the HSP90 inhibitor luminespib and the mTORC1/2 inhibitor inhibitor torin-2 induced strong cytotoxicity in most KRAS WT PDOs. Together, these observations reflect the dependency of KRAS-WT PDOs on upstream receptor-driven signaling, and their marked susceptibility to downstream pathway collapse when proteostasis or PI3K/mTOR is disrupted. In parallel, cytostatic responses were predominantly enriched among RTK inhibitors, including afatinib (irreversible ErbB family blocker), lapatinib (dual EGFR/HER2 inhibitor) and brivanib (VEGFR/FGFR inhibitor) as well as the WNT signaling inhibitor LGK-974. Across the cohort, numerous compounds targeting key oncogenic programs including, cell-cycle regulation, MAPK signaling, the DNA damage response, RTK pathways, and TGF-β signaling, elicited effective cytostatic responses in distinct PDOs, indicating that growth arrest is a common therapeutic outcome across genetically diverse tumors. This pattern suggests that, in KRAS WT backgrounds, proliferative signaling can be effectively arrested without immediate loss of structural integrity.

In contrast, within the KRAS-mutant PDO subset, fewer agents induced cytotoxicity, consistent with the intrinsically resilient and adaptive signaling architecture of these PDOs **(Fig. 3b)**. Nevertheless, notable vulnerabilities were uncovered. For example, similar to the KRAS-WT cohort, bortezomib, luminespib, and torin-2 remained potent cytotoxic agents in all three KRAS-mutant PDOs, while the MEK1/2 inhibitor trametinib also emerged as a cytotoxic hit, indicating that direct MEK blockade can override adaptive MAPK signaling even in KRAS-mutant PDOs. Furthermore, most cytotoxic and cytostatic hits in KRAS-mutant PDOs targeted key regulators of the cell-cycle and DNA damage response, including rigosertib (RAS-mimetic disrupting RAS–microtubule interactions), adavosertib (WEE1 inhibition driving mitotic entry under replication stress), AZD7762 (CHK1/2 inhibitor impairing checkpoint activation), danusertib (pan-Aurora kinase blockade disrupting spindle assembly), and palbociclib (CDK4/6 inhibitor preventing G1–S transition). The recurrent sensitivity of these KRAS-mutant PDOs to agents that perturb replication surveillance and mitotic control underscores a shared dependency on checkpoint integrity as an adaptive survival mechanism downstream of oncogenic KRAS, potentially reflecting the elevated proliferative pressure characteristic of these models. Collectively, These observations align with the studies demonstrating that KRAS-mutant CRC organoids frequently acquire adaptive resistance via compensatory pathways rewiring, thereby highlighting the need for combinatorial targeting of both RAS–MAPK cascade and DNA damage response or cell-cycle regulatory modules^32,34^. Given that KRAS-mutant PDOs predominantly displayed cytostatic overt cytotoxicity, our data strongly support the prioritization of rational combination strategies, integrating cytostatic and cytotoxic agents, over single-agent therapies for this genetically defined subgroup. To illustrate these divergent response patterns, representative maximum-projection images of the most effective cytotoxic and cytostatic compounds are shown for selected PDOs **(Fig. 3c)**. Notably, these agents did not elicit uniform efficacy across all models, highlighting the pronounced inter-patient heterogeneity that characterizes therapeutic response in metastatic CRC. Collectively, our multi-parametric, imaging-based drug screening approach not only maps inter-patient heterogeneity in drug response but also uncovers a clear association between KRAS mutational status and the nature of response (cytotoxic vs cytostatic). When integrated with single-organoid-resolved phenotypic profiling, these data establish a refined functional stratification framework to guide the rational selection of therapeutic backbones in metastatic CRC.

### Clonal and molecular characterization of BI-2865 resistance in KRAS-WT and KRAS-Mutant PDOs

Building on the genomic and transcriptomic characterization of our PDO biobank, we next focused on two representative PDO models to investigate the mechanisms of acquired resistance to the novel pan-KRAS inhibitor BI-2865. This transition from cohort level profiling to in-depth functional modeling enabled controlled dissection of therapy-induced resistance at the clonal and molecular levels. For this purpose, PDOs with high proliferative capacity, PDO-1 (KRAS ^WT^) and PDO-6 (KRAS^G13D^), were selected for long-term drug exposure studies. To enable tracking of clonal dynamics during resistance evolution, these PDOs were transduced with a high-complexity Cellecta® lentiviral expressible barcode library containing approximately one million unique barcodes. Each barcode was designed to generate a polyadenylated transcript compatible with capture by single-cell RNA sequencing. To ensure that each cell received a single unique barcode, transductions were performed at a low multiplicity of infection (M.O.I.) of ∼0.1 **(Supplementary Fig. 2**). Following barcode labelling, PDO populations were divided into four experimental arms per model. Three independent replicates were subjected to continuous exposure of the non-covalent pan-KRAS inhibitor BI-2865, while matched DMSO-treated controls were maintained in parallel **(Fig. 4a)**. Resistance was induced through stepwise dose escalation over a three-month treatment period, starting from IC50/10 and increasing the drug concentration at each medium change until reaching 2×IC50, resulting in the emergence of drug-tolerant and resistant populations. At the end of this evolutionary selection period, dose-response analyses demonstrated that replicates A, B, and C from both PDO-01 and PDO-06 had acquired resistance relative to DMSO-treated controls, pre-resistant baseline populations, and unbarcoded parental counterparts **(Fig. 4b and Fig. 4c)**.

**Figure 4.**
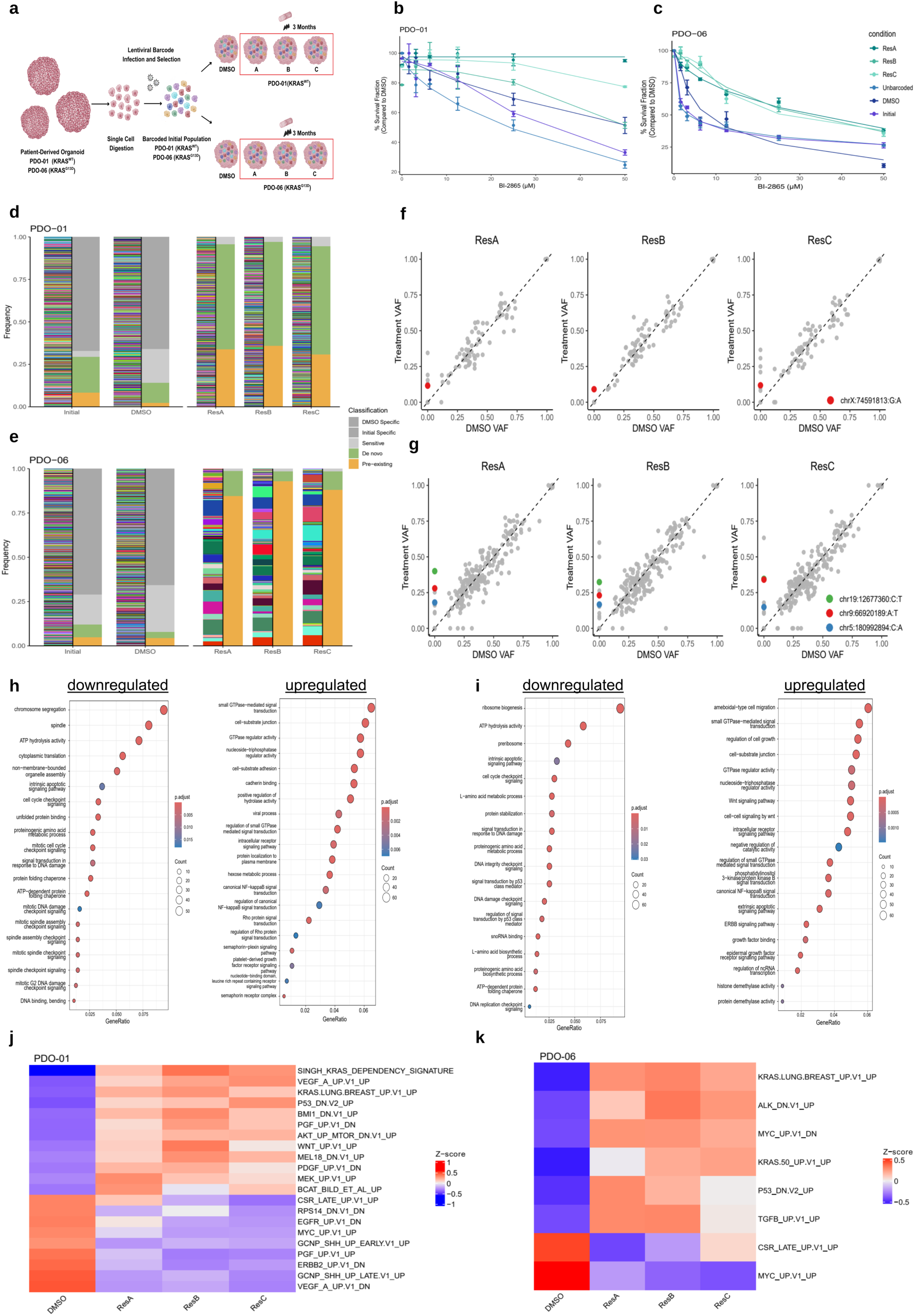
Lineage tracing and molecular characterization of BI-2865 resistance in patient-derived organoid models. **a** Experimental strategy for tracing clonal evolution during acquisition of BI-2865 resistance. Two highly proliferative PDO models, PDO-01 (KRAS^WT^) and PDO-06 (KRAS^G13D^), were transduced at low multiplicity (MOI = 0.1) with a Cellecta® lentiviral barcode library (∼1 million poly-A–tagged barcodes) to enable lineage tracking and transcript capture. Barcoded populations were expanded and split into four arms: three replicates subjected to long-term BI-2865 treatment and a matched DMSO control. Resistant populations were established by stepwise dose escalation over three months. **b&c** Dose–response analyses demonstrating acquired BI-2865 resistance. PDO-01 (KRAS^WT^ panel b) and PDO-06 (KRAS^G13D^ panel c) were exposed to increasing BI-2865 concentrations, and viability was measured relative to DMSO controls using CellTiter-Glo 3D. Curves are shown for the initial barcoded population (Initial), an unbarcoded control (Unb), the barcoded DMSO control, and three independently evolved resistant replicates (ResA–C). Resistant populations exhibit increased BI-2865 IC₅₀ values compared to controls (mean ± s.e.m., n = 3). **d&e** Barcode frequency distributions during BI-2865 selection. Clonal composition was assessed in PDO-01 (panel d) and PDO-06 (panel e) at baseline (Initial), in barcoded DMSO controls, and in three BI-2865-resistant replicates. Individual barcodes were classified as pre-existing, de novo, or sensitive based on comparative frequency changes, revealing treatment-associated shifts in clonal representation across replicates. **f&g** Variant allele frequency (VAF) analysis of BI-2865 selection. Scatter plots compare VAFs from WES of resistant replicates versus matched DMSO controls for PDO-01 (panel f) and PDO-06 (panel g). Each point represents a somatic variant; deviations from the diagonal indicate altered clonal representation following BI-2865 exposure. **h&i** Gene ontology enrichment in BI-2865-resistant PDOs relative to DMSO controls. Gene Ontology Enrichment Analysis of DEGs identifies pathways altered in PDO-01 (panel h) and PDO-06 (panel i), with downregulated pathways shown on the left and upregulated pathways on the right. Bubble size denotes gene counts and color indicates adjusted p-values; results are shown across three resistant replicates. **j&k** Oncogenic C6 signature enrichment in BI-2865-resistant PDOs. Heatmaps display GSVA-derived C6 oncogenic signature scores for PDO-01 (panel j) and PDO-06 (panel k) across DMSO controls and resistant replicates (ResA–C). Values are shown as z-scores, with red and blue indicating relative enrichment or depletion.

Integration of high-complexity cellular barcoding with BI-2865 treatment provided a powerful experimental framework to dissect how clonal heterogeneity and evolutionary pressure jointly shape acquired resistance in PDOs. Barcode tracking revealed divergent patterns of clonal evolution between KRAS wild-type and KRAS mutant PDOs under BI-2865 treatment. In the KRAS wild-type PDO-01 model, resistance predominantly emerged through the *de novo* outgrowth of previously drug-sensitive clones, indicative of adaptive resistance arising under therapeutic selective pressure **(Fig. 4d)**. This evolutionary pattern is consistent with models of drug-induced plasticity, in which non-genetic state transitions contribute to resistance acquisition^35^. In contrast, in the KRAS G13D-mutant PDO-06 model, resistance was driven primarily by the selective expansion of pre-existing resistant clones, reflecting a typical clonal selection process during prolonged drug exposure **(Fig. 4e)**. This finding indicates the presence of intrinsically drug resistant subpopulations prior to treatment, plausibly facilitated by the KRAS mutation^36^. Collectively, these findings delineate two distinct Darwinian modes of therapeutic escape, adaptive plasticity in KRAS wild-type PDOs versus intrinsic clonal resistance in KRAS-mutant PDOs, underscoring the genotype-dependent nature of acquired resistance in metastatic colorectal cancer.

To determine whether clonal shifts were associated with underlying genetic or transcriptional adaptations, we next performed WES and bulk RNA sequencing on matched resistant and control populations from both PDO models. WES analysis of pan-KRAS inhibitor–resistant replicates revealed that the global mutational landscapes remained largely conserved relative to DMSO-treated controls, indicating that resistance did not arise through widespread genomic remodeling. However, a small subset of variants exhibited consistent increases in variant allele frequencies (VAF) following resistance acquisition, suggesting positive selection of specific genetic alterations under therapeutic pressure. In the KRAS wild-type PDO-1, resistance was reproducibly associated with enrichment of a recurrent missense variant in *RNF12* (chrX:74591813 G>A) across independent replicates **(Fig. 4f)**. In the KRAS G13D PDO-6 model, resistance was associated with increased VAFs of missense variants in *DHPS* (chr19:12677360 C>T), *ZNF658* (chr9:66920189 A>T) and *BTNL3* (chr5:180992894 C>A) **(Fig. 4g)**. Although these genes operate in distinct molecular pathways, they converge functionally on cellular stress adaptation and proliferative fitness. RNF12 modulates the stability of key stress-response and cell-cycle regulators, including p53 and c-Myc^37–39^. DHPS, which catalyzes hypusination of eIF5A, is recurrently overexpressed in multiple cancer types and associated with aggressive disease, poor prognosis and, in some settings, resistance to therapy; notably, pharmacological or genetic inhibition of DHPS. For example, next-generation DHPS inhibitors can suppress tumor growth and restore drug sensitivity in preclinical models^40–42^. ZNF658 belongs to a zinc-finger transcriptional repressor family linked to stress-responsive transcriptional remodeling^43^. BTNL3, a butyrophilin-like molecule, participates in the regulation of γδ T-cell responses at epithelial barriers, and BT(N/L) family members have emerged as immunoregulatory checkpoints in solid tumours, including colorectal cancer, where their dysregulation can facilitate immune escape and modulate responses to a therapy^44–47^. Together, these findings suggest that pan-KRAS inhibitor pressure selectively enriches subclones harboring genetic alterations that enhance stress tolerance, proteostatic control, and epithelial immune adaptability, thereby providing a genomic foothold for the emergence and stabilization of acquired resistance.

Transcriptomic profiling of BI-2865-resistant KRAS wild-type PDO-01 replicates revealed a coordinated suppression of programs governing cell-cycle progression, DNA-damage checkpoint activity, apoptotic signaling, translational machinery, and amino-acid biosynthesis, reflected by consistent downregulation of corresponding hallmark pathways **(Fig. 4h)**. This transcriptional pattern suggests that resistant cells transition into a slow cycling, metabolically restrained, drug-tolerant state, a phenotype arguably associated with cost of resistance and that has been described in in MAPK pathway inhibited cancers^48–50^ . In parallel, resistant PDOs displayed coordinated upregulation of pathways associated with NF-κB activation, Rho/small-GTPase-driven cytoskeletal remodeling, cell–substrate adhesion, cadherin-mediated junctions, semaphorin–plexin and PDGFR-associated signaling. This transcriptional reprogramming is consistent with acquisition of an adhesion anchored, mechanically adaptive survival state, in which NF-κB signaling provides anti-apoptotic support while adhesion and cytoskeletal programs enhance cellular persistence, characteristics of drug tolerant persister phenotypes under MAPK suppression^51–54^. The concomitant induction of semaphorin–plexin and PDGF-axis signatures further suggests engagement of MAPK-independent pro-survival signals that can sustain cell viability despite KRAS pathway inhibition^55,56^. Analysis of oncogenic C6 signatures further supported this transcriptional pattern **(Fig. 4j)**. Across all BI-28650-resistant replicates, robust enrichment of KRAS dependency gene sets (*SINGH_KRAS_DEPENDENCY, KRAS.LUNG.BREAST_UP*) was observed, indicating that resistant cells preserve core elements of a KRAS-driven transcriptional program despite a marked suppression of proliferative output. Consistent enrichment of MEK_UP and AKT/mTOR-associated signatures further suggests that intracellular MAPK and PI3K/AKT/mTOR modules remain transcriptionally active, providing a compensatory signaling backbone capable of sustaining viability under KRAS inhibition. Similarly, enrichment of WNT/β-catenin-associated signatures (*WNT_UP; BCAT_BILD_ET_AL_UP*) suggests that the activation of epithelial survival programs which is frequently engaged when MAPK activity is attenuated^57^. In contrast, several receptor tyrosine kinase-related oncogenic signatures including, *EGFR_UP.V1_DN*, *ERBB2_UP.V1_DN*, and *VEGF_A_UP.V1_DN* were consistently downregulated across replicates, indicating a functional shift away from upstream RTK-mediated growth signaling. Together, this coordinated suppression of EGFR/ERBB/VEGF signaling, alongside with preserved MAPK/AKT/WNT axis activity, supports a model in which resistant KRAS-WT organoids become progressively less dependent on extrinsic growth factor signaling and instead stabilize an intracellular, KRAS/MAPK-dependent survival strategy that enable long-term persistence under therapeutic blockade.

Transcriptomic profiling of BI2865-resistant KRASG13D PDOs revealed a global adaptive program that, while partially overlapping with the KRAS-WT model, exhibited distinct resistance-associated gene expression profile. Similar to BI2865-resistant resistant KRAS wild-type PDO-01, BI2865-resistant KRASG13D PDO-06 replicates exhibited consistent suppression pathways governing cell-cycle checkpoints, DNA-damage response, intrinsic apoptosis, ribosome biogenesis, protein synthesis, and amino-acid metabolism **(Fig. 4i)**. The concurrent attenuation of p53-mediated signaling further aligns with established mechanisms by which cancer cells evade therapy by weakening p53-dependent apoptotic surveillance^58^. In contrast to this quiescence-associated program, resistant KRAS G13D PDOs showed reproducible activation of pathways related to small GTPase/Rho signaling, WNT signaling and WNT-driven self-renewal, PI3K/AKT signaling, canonical NF-κB activation, ERBB/EGFR signaling, and cell-substrate junction organization. This coordinated transcriptional state reflects multiple cooperating resistance mechanisms: Rho GTPase-driven cytoskeletal remodeling supports adaptive motility and treatment evasion^59^; WNT/β-catenin activation promotes self-renewal and therapy resistance in colorectal cancer ^60^; and PI3K/AKT together with NF-κB confer potent anti-apoptotic and pro-survival advantages in KRAS-driven epithelial tumors^61^. Notably, upregulation of ERBB/EGFR signaling suggests activation of compensatory RTK signaling, a well-described resistance mechanisms in KRAS-mutant colorectal cancer following MAPK pathway supression^62^. Together, these data indicate that KRAS G13D–driven resistance to pan-KRAS inhibition is sustained through the simultaneous engagement of cytoskeletal plasticity, stemness-associated WNT programs, pro-survival PI3K/AKT–NF-κB signalling, and compensatory ERBB/EGFR activation, thereby establishing a highly resilient, multi-layered survival state under therapeutic pressure.

Oncogenic C6 signatures further supported the resistant transcriptional state of KRAS G13D PDOs **(Fig. 4k)**. Consistent enrichment of *KRAS.LUNG.BREAST_UP* and *KRAS.50_UP* confirmed that KRAS-linked transcriptional programs remain active in the resistant setting. In parallel, the upregulation of *TGFB_UP* and *CSR_LATE_UP* signatures indicated a shift toward TGF-β driven cellular plasticity and a serum-response-like wound-healing program, both of which are known to promote motility and stress adaptation^63,64^. Elevated *P53_DN* signatures were consistent with weakened apoptotic surveillance in the resistant state. Notably, MYC-associated signatures showed a bidirectional pattern (*MYC_UP.V1_UP* ↑, *MYC_UP.V1_DN* ↓), suggesting selective retention of MYC-linked metabolic and stress-adaptive outputs alongside suppression of MYC-driven proliferative programs. Collectively, these features define a highly plastic, growth factor responsive KRAS G13D resistance state that is mechanistically distinct from the attenuated growth and mitogenic signaling, adhesion anchored, metabolically restrained phenotype observed in the KRAS-WT model. Only significantly altered C6 signatures are displayed for each PDO model.

### Resistance-Driven Functional Vulnerabilities in KRAS-Wild-Type and KRAS-Mutant CRC PDOs

As previously described that 3D image-based high-content drug screening was performed with a focused objective to identify therapeutic vulnerabilities that specifically emerge following the acquisition of BI-2865 resistance in KRAS-WT and KRAS-Mutant PDOs. In this context, our analysis focused on cytotoxic hits, defined as compounds that reduced the viability of resistant PDOs more strongly than their treatment-naïve counterparts. This approach enabled the systematic detection of resistance associated vulnerabilities. Drug screening heatmaps revealed striking, genotype-dependent therapeutic vulnerabilities that emerge after the BI-2865 resistance **(Fig. 5a)**, highlighting that resistance reshapes the spectrum of actionable drug responses. In the KRAS-WT model (PDO-01), resistant PDOs acquired selective sensitivity to cell cycle and checkpoint inhibitors, including Chk1/2 (AZD7762), WEE1 (Adavosertib), CDK4/6 (Palbociclib) and Aurora kinase inhibitors (Barasertib, ENMD-2076). These dependencies are consistent with the RNA-seq analysis results of cell cycle suppression, attenuated p53 signaling and enhanced checkpoint dependence. In parallel, newly acquired vulnerabilities to mTOR/PI3K inhibitors (Torin, Apitolisib, AZD6482) and ERBB/RTK inhibitors (Dacomitinib, Gefitinib) indicate that pan-KRAS inhibitor-resistant WT PDOs retain a minimal but targetable dependence on RTK–PI3K–mTOR signaling for survival within their adhesion-anchored, persister like state. Collectively, these patterns show that resistance in the KRAS-WT genotype generates a checkpoint-fragile phenotype, exposing functionally actionable vulnerabilities that are not present in the drug-sensitive state.

**Figure 5.**
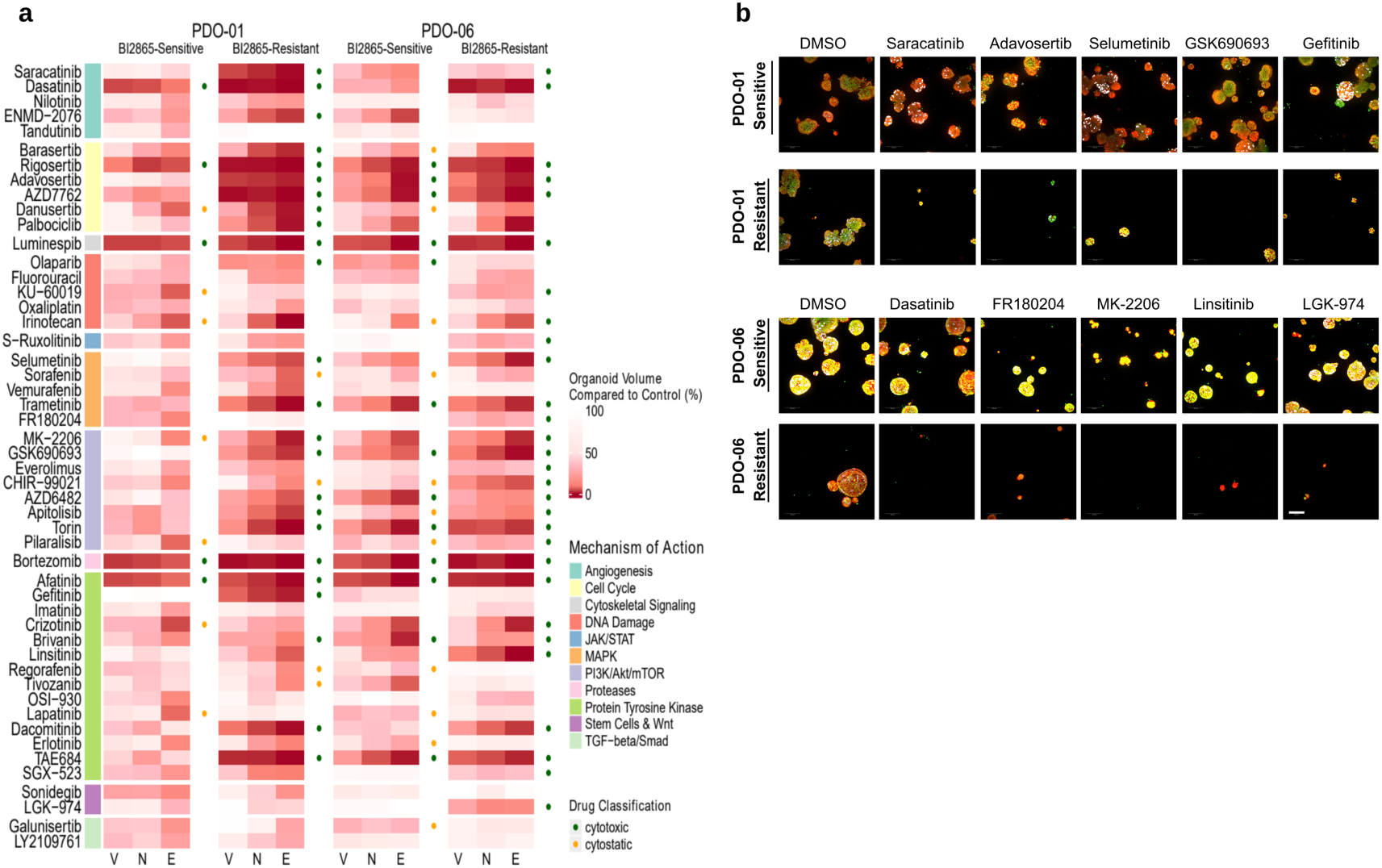
Functional drug vulnerability profiling of BI-2865-resistant patient-derived organoids. **a** Phenotypic drug screening of BI-2865–sensitive and BI-2865–resistant PDOs. PDO-01 (KRAS^WT^) and PDO-06 (KRAS^G13D^) were cultured for 4 days to allow organoid formation, followed by a 6-day single-dose treatment (1 µM), yielding a 10-day assay. A panel of 50 targeted compounds was profiled using high-content 3D imaging, and responses were quantified by organoid volume (V), total nuclei count (N), and EdU-positive nuclei fraction (E). Heatmaps show percent change relative to DMSO controls for sensitive and resistant populations. Compounds were classified as cytotoxic (green dots) when V, N, and E were jointly reduced or when combined loss of V and N exceeded 50%, and as cytostatic (yellow dots) when structural integrity was preserved (V and N > 50%) despite strong suppression of proliferation (E < 50%). **b** Representative 3D images of drug responses in BI-2865–sensitive versus resistant PDOs. Maximum-intensity projections illustrate phenotypic effects of selected compounds targeting distinct pathways, highlighting differential cytotoxic responses in resistant populations. Confocal Z-stacks were acquired on an Operetta CLS system (20× water-immersion objective, NA = 1.0) across five fields per well with 1 µm Z-steps. Scale bar, 100 µm.

In contrast, the KRAS G13D-mutant model (PDO-06) exhibited a broader and more pronounced vulnerability landscape **(Fig. 5a)**, consistent with its transcriptional shift toward a signal-amplified, growth factor-responsive resistant state. BI-2865-resistant KRAS G13D PDOs acquired selective sensitivity to inhibitors targeting RTKs (Crizotinib, Linsitinib, SGX-523), SRC-family kinases (Dasatinib, Saracatinib), MEK/ERK signaling (Selumetinib, FR180204) and the PI3K/AKT/mTOR axis (MK-2206, Everolimus). These functional dependencies mirror the activation of Rho-GTPase, WNT, PI3K/AKT, NF-κB and ERBB/EGFR programs identified by RNA sequencing. Notably, the most striking newly acquired dependency was a potent and selective vulnerability to PORCN inhibition (LGK-974), a hit that was absent in the treatment-naive state, directly concordant with the WNT/β-catenin pathway upregulation unique to the KRAS G13D resistance program. This finding suggests that BI-2865 resistance in the KRAS G13D context might drive a WNT-ligand-dependent survival architecture, converting WNT secretion into a druggable vulnerability. Together, these functional screening patterns demonstrate that post-resistance drug screening not only identifies resistance-specific therapeutic vulnerabilities but also provides validation of the molecular rewiring responsible for BI-2865 treatment escape. KRAS-WT-resistant PDOs reveal vulnerabilities rooted in checkpoint fragility and residual RTK-PI3K support, whereas G13D-resistant organoids expose dependencies driven by amplified RTK-WNT-AKT signaling and cellular plasticity, highlighting mechanistically distinct and therapeutically actionable routes to overcome pan-KRAS inhibitor resistance. Finally, to further substantiate these resistance-specific dependencies at the phenotypic level, we included representative high-content imaging of five mechanistically distinct inhibitors that induce a cytotoxic collapse exclusively in resistant PDOs while remaining largely ineffective in their treatment-naïve counterparts. These images provide PDO-based visual confirmation of the acquired resistance vulnerabilities defined by the screen, illustrating the marked structural disintegration observed only under post-resistance conditions **(Fig. 5b)**.

### Single-Cell and Lineage Tracing Reveal Clonal Redistribution in the BI-2865 Resistant KRASG13D Mutant PDO Model

Among the two genotypes analyzed by bulk omics, BI-2865-resistant KRAS^G13D^ PDOs showed a markedly broader adaptive signaling landscape and pronounced clonal selection dynamics, suggesting that resistance arises through heterogeneous trajectories rather than a uniform shift in cell-state. For this reason, we next focused on the KRAS^G13D^ PDO model to resolve the cellular programs underlying KRAS inhibitor resistance at single-cell resolution. Importantly, with the use of expressible lentiviral barcoding approach, we sought to distinct clonal lineages under BI-2865 pressure. To resolve the cellular architecture of KRAS inhibitor resistance at single-cell resolution, we profiled BI-2865–resistant KRAS^G13D^ organoids and matched DMSO controls using scRNA-seq combined with lentiviral expressible barcoding. UMAP embedding of ∼35000 cells demonstrated that resistant and control populations occupy a partially shared transcriptional program **(Fig. 6a)**, indicating that acquired resistance does not involve the emergence of a novel drug-induced cell state. Instead, resistant cells showed partial spatial divergence within this common landscape, forming localized enrichments in discrete regions of the existing state-space rather than separating into a totally independent cluster. This pattern suggests that BI-2865 resistance arises through selective re-weighting of pre-existing phenotypic variation, rather than wholesale transcriptional reprogramming. In addition, cell-cycle phase assignment revealed comparable distributions of G1, S and G2/M states between control and resistant cells **(Fig. 6b)**, indicating that the resistant phenotype is maintained without a dominant shift in proliferative status at the population level. Cluster resolved analyses further showed modest, state-specific differences in cell-cycle phase distribution across conditions, without evidence of a global reprogramming of cell-cycle activity **(Supplementary Fig. 3).**

**Figure. 6.**
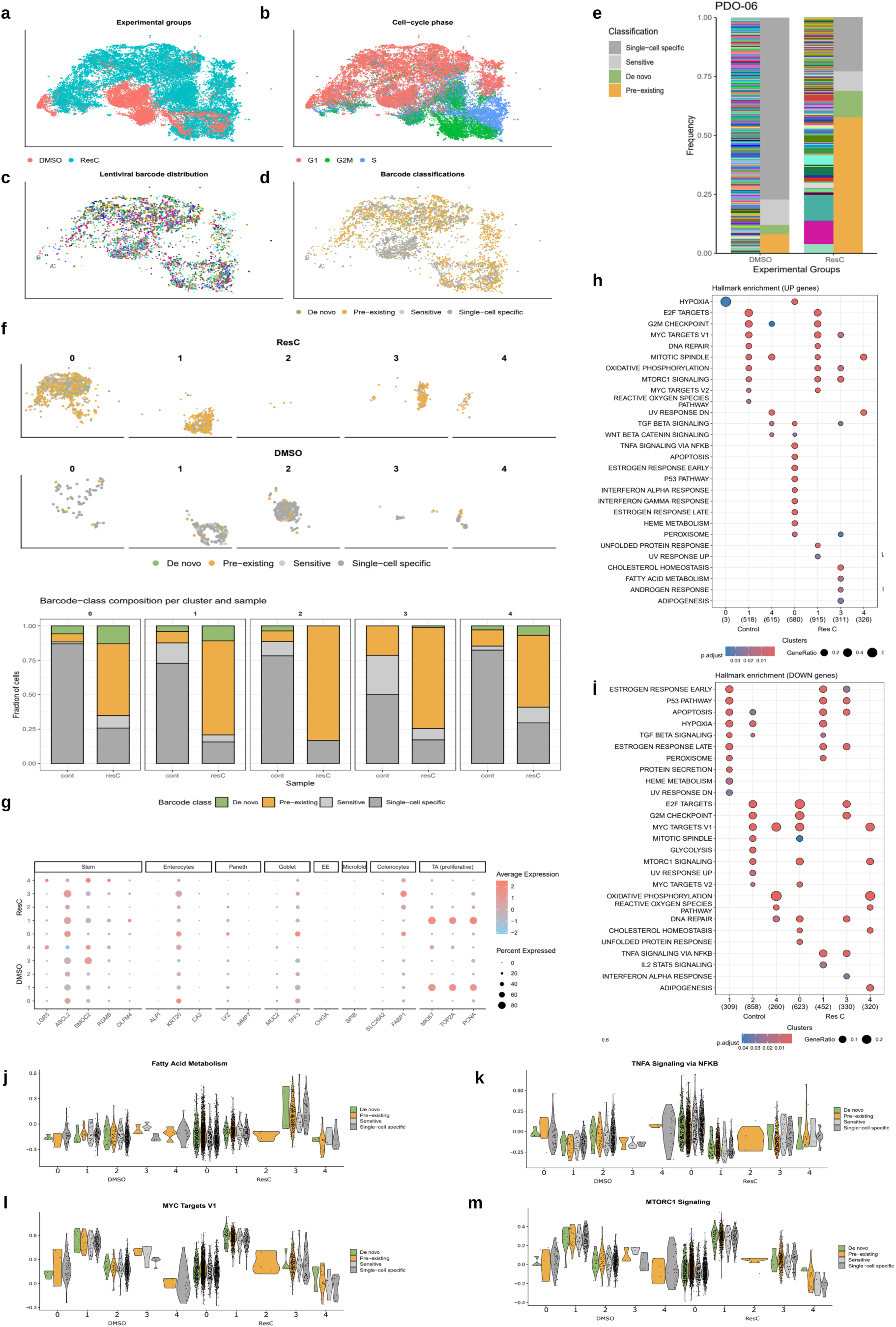
Single-cell transcriptomics reveals heterogeneous clonal trajectories underlying KRAS inhibitor resistance in KRAS^G13D^ organoids. **a** UMAP embedding of single-cell transcriptomes from DMSO-treated controls and BI-2865–resistant cells (replicate C) show a continuous transcriptional manifold without evident discrete treatment-specific clusters. **b** Cell-cycle phase mapping onto the same UMAP indicates comparable distributions of G1, S, and G2/M phases between control and resistant cells. **c** Projection of expressible lentiviral barcode identities onto the UMAP reveals extensive intermixing of barcoded lineages across shared transcriptional states. **d** Clonal fate assignment (pre-existing, de novo, sensitive) was performed by calling barcode identities at the DNA level and matching them to expressible barcode transcripts detected in single-cell RNA-seq. Cells lacking a resolvable barcode class due to resolution limitations of DNA barcode sequencing technology were labeled as single-cell specific due to absence of lentiviral integration or barcode transcript capture, are shown separately and excluded from fate categories. **e** Barcode lineage frequency distribution in PDO-06. Stacked bar plots show relative frequencies of barcoded lineages in DMSO-treated controls and BI-2865–resistant populations. Pre-existing and de novo lineages expand under drug pressure, whereas sensitive and single cell specific barcodes are reduced; NA cells are displayed separately. **f** Cluster-based analysis of clonal fate. UMAP projections and stacked bar plots show distribution of barcode classes across five transcriptional clusters (0–4) in control and resistant samples, demonstrating cluster-dependent shifts from sensitive to pre-existing and de novo lineages**. g** Intestinal lineage marker expression across clusters in control and BI-2865–resistant PDO-06. Dot size indicates the fraction of expressing cells per cluster, and color denotes z-scored average expression, showing preservation of major epithelial lineages across transcriptional states. **h&i** Cluster-resolved Hallmark pathway enrichment across conditions. Dot plots showing Hallmark pathway enrichment of upregulated (panel h) and downregulated (panel i) genes across transcriptional clusters in DMSO control and resistant (ResC) conditions. Dot size represents gene ratio and color indicates adjusted p value, highlighting state-specific pathway activation and suppression patterns associated with resistance. **j&m** Violin plots of single-cell pathway activity scores for fatty acid metabolism (panel j), TNFα signaling via NF-κB (panel k), MYC targets (V1) (panel l), and mTORC1 signaling (panel m) are shown across transcriptional clusters under DMSO control and resistant (ResC) conditions. Violin plots depict the distribution of pathway activity at single-cell resolution, stratified by clonal fate classes.

Expressible barcodes mapped onto the UMAP revealed extensive lineage mixing across shared cell-state space **(Fig. 6c)**, confirming that multiple barcoded clones can occupy overlapping transcriptional regions. Barcode classes were derived by matching DNA-level barcode identities to barcode sequences detected in scRNA-seq reads **(Fig. 6d)**. As scRNA-seq was generated from one resistant replicate, some barcodes present in the DNA library did not have detectable transcript reads and were labeled single-cell specific, but were retained in downstream analyses to maintain clonal diversity and avoid expression bias; notably, single-cell specific barcodes showed no preferential enrichment within specific regions of the manifold, indicating that their absence reflects detection limitations rather than a distinct biological state. Notably, single-cell specific barcodes were abundant, particularly in the DMSO control condition, consistent with the higher sensitivity and depth of scRNA-seq in capturing low-frequency or transient barcodes at the single-cell level. Classification of valid barcode fates showed that pre-existing resistant clones are broadly distributed across the manifold, consistent with high phenotypic plasticity and multiple pre-treatment states supporting survival. Consistent with DNA barcode results, single-cell barcode frequency analysis also demonstrated that pre-existing lineages constitute the largest clonal fraction under BI-2865 treatment, markedly exceeding the proportion of de novo lineages **(Fig. 6e)**, further supporting a dominant contribution of pre-existing resistance reservoirs. By contrast, de novo resistant clones form localized enrichments within defined subregions, indicating that new adaptions emerge through constrained consolidation of survival programs rather than acquisition of a distinct cell identities. Thus, acquired BI-2865 resistance in KRAS^G13D^ occurs through divergent clonal trajectories within a shared transcriptional landscape, rather than through the formation of a new state **(Fig. 6d)**. Together, these data show that acquired resistance in KRAS^G13D^organoids is not governed by the creation of a new transcriptional identity, but rather by redistribution of clonal trajectories within a shared phenotypic landscape, enabling distinct evolutionary solutions to emerge from pre-existing cell-state space. This architecture points to multiple adaptive pathways embedded within the G13D genetic background.

To resolve whether distinct resistant cell states arise from different clonal origins, we quantified barcode-class composition within the five transcriptional clusters defined by scRNA-seq **(Fig. 6f)**. All resistant clusters contained resistant cells, but their clonal architecture separated into two clear patterns. Clusters 2 and 3 were almost entirely composed of pre-existing resistant lineages, indicating deep selection of specific pre-treatment clones that can support long-term survival without requiring new adaptive events. By contrast, Clusters 0, 1 and 4 showed dominant pre-existing contributions together with a consistent fraction of de novo lineages, reflecting mixed adaptive solutions in which new resistance trajectories emerge alongside pre-existing reservoirs. Sensitive barcodes persisted but at substantially lower frequencies across clusters under BI-2865 treatment. Together, these data demonstrate that acquired BI-2865-resistance in KRAS^G13D^ is supported by multiple cluster-specific survival states; some seeded entirely from pre-existing diversity, and others involving limited de novo contributions within defined transcriptional neighborhoods.

To determine whether resistance-associated clusters adopt distinct epithelial identities, we profiled colon lineage markers across clusters in control and BI-2865–resistant organoids **(Fig. 6g)**. Across all resistant clusters, we observed upregulation of regenerative stem-like markers, including *RGMB* and *OLFM4*, indicating that BI-2865 resistance is accompanied by a shift toward a crypt-base stem identity. In parallel, goblet differentiation marker *MUC2* was reduced, consistent with suppression of the secretory lineage and bias toward a regenerative program. Notably, *FABP1* was selectively enriched in resistant clusters, together with increased expression of additional absorptive colonocyte markers, suggesting a metabolic adaptation toward lipid handling within the resistant state. TA/proliferative markers (*MKI67*, *TOP2A*, *PCNA*) were largely maintained rather than expanded, indicating that resistance is driven by lineage remodeling rather than hyper-proliferation. Taken together, these data reveal that KRAS^G13D^ resistance involves coordinated epithelial plasticity, in which a WNT-driven regenerative identity and FABP1-associated metabolic reprogramming emerge while secretory differentiation is suppressed.

To further dissect how these lineage and metabolic features are distributed across resistant cell states, we next performed cluster-resolved pathway analysis comparing control and BI-2865–resistant PDO. Cluster-resolved Hallmark pathway analysis revealed that BI-2865 resistance in KRAS^G13D^ PDO is associated with heterogeneous redistribution of functional programs rather than uniform pathway activation across the resistant population. Several pathways were similarly enriched in both control and resistant conditions within specific clusters, indicating stable, cluster-intrinsic transcriptional states. For example, hypoxia signaling was comparably upregulated in both control and resistant cells within Cluster 0, while Cluster 1 consistently displayed enrichment of proliferative and cell-cycle–associated programs, including MYC TARGETS (V1 and V2), MTORC1 SIGNALING, OXIDATIVE PHOSPHORYLATION, MITOTIC SPINDLE, DNA REPAIR, G2M CHECKPOINT, and E2F TARGETS, irrespective of treatment condition. These shared signatures indicate that canonical MYC–cell cycle programs represent baseline features of specific cellular states rather than resistance-induced signaling events **(Fig. 6h&i).**

In contrast, resistant clusters exhibited selective, cluster-specific enrichment of distinct pathway modules that were not observed in their matched control counterparts. Resistant Cluster 0 showed preferential upregulation of plasticity and stress-associated programs, including WNT/β-CATENIN SIGNALING, TGF-BETA SIGNALING, TNFA SIGNALING VIA NFκB, INTERFERON ALPHA and GAMMA RESPONSES, as well as HEME METABOLISM and PEROXISOME pathways, accompanied by relative attenuation of proliferative programs such as MYC TARGETS, MTORC1 SIGNALING, G2M CHECKPOINT, E2F TARGETS, and DNA REPAIR. Resistant Cluster 1 was characterized by selective enrichment of stress-tolerance pathways, most prominently UNFOLDED PROTEIN RESPONSE and UV RESPONSE UP, consistent with a proteostasis and damage adaptive transcriptional state. By contrast, Resistant Cluster 3 displayed coordinated upregulation of metabolically reinforced programs, including OXIDATIVE PHOSPHORYLATION, MTORC1 SIGNALING, FATTY ACID METABOLISM, CHOLESTEROL HOMEOSTASIS, ADIPOGENESIS, PEROXISOME, and ANDROGEN RESPONSE, alongside MYC TARGETS V1, while inflammatory and damage-response pathways such as INTERFERON ALPHA RESPONSE, TNFA SIGNALING VIA NFκB, APOPTOSIS, P53 PATHWAY, and cell-cycle checkpoint programs were comparatively attenuated **(Fig. 6 h&i)**.

Importantly, this cluster-specific pattern resulted in several pathways appearing upregulated in one resistant cluster while being downregulated in another, reflecting functional partitioning of proliferative, metabolic, plasticity, and stress-response modules across transcriptional neighborhoods within a shared cellular state space. Together, these findings indicate that KRAS^G13D^ resistance emerges through differential deployment and redistribution of pre-existing cellular programs rather than through induction of a single, uniform resistance-associated signaling pathway. This modular organization of resistance-associated programs is consistent with prior studies indicating that therapeutic resistance often reflects redistribution of pre-existing transcriptional states rather than induction of a single resistance-defining pathway^65,66^.

To further resolve how resistance-associated programs are distributed across resistant cell states, we examined the cluster-resolved activity of selected Hallmark pathways at single-cell resolution. Fatty acid metabolism showed selective enrichment in resistant Cluster 3, where higher pathway scores were observed across clonal classes, indicating concentration of lipid metabolic programs within a metabolically specialized transcriptional niche rather than global activation across resistant cells **(Fig. 6j)**. In contrast, TNFA signaling via NFκB displayed opposing patterns across clusters, with relative enrichment in resistant Cluster 0 and attenuation in Cluster1 and 3, highlighting context-dependent deployment of inflammatory stress-response programs **(Fig. 6k)**. MYC Targets V1 activity remained high in Cluster 1 under both control and resistant conditions and was broadly distributed across clonal classes, consistent with a cluster-intrinsic proliferative state that is maintained rather than induced upon resistance **(Fig. 6l)**. Similarly, MTORC1 signaling showed a cluster-dependent pattern, with higher activity in metabolically reinforced clusters and lower activity in others, indicating context-dependent utilization rather than uniform activation upon resistance **(Fig. 6m)**. Together, these pathway-specific distributions demonstrate that resistance-associated signaling is organized in a modular, cluster-dependent manner, with distinct metabolic, proliferative, and inflammatory programs preferentially localized to defined cellular neighborhoods within a shared transcriptional landscape.

## Discussion

In this study, we report the establishment of 20 mCRC PDOs and a comprehensive experimental framework that integrates PDOs, multiparametric drug phenotyping, and lineage-resolved molecular profiling to elucidate the evolutionary mechanisms driving therapeutic resistance in metastatic CRC. PDOs serve as physiologically relevant ex vivo models that preserve the genomic architecture and phenotypic complexity of patients’ tumors, thereby capturing the inter- and intra-tumoral heterogeneity that shapes clinical treatment response. In line with previous large-scale organoid studies^9,10,24,67^, our mCRC PDOs maintained the mutational and transcriptional hallmarks of their parental tumors, validating their utility as functional avatars for precision drug testing. This model system not only reflects static tumor characteristics but also enables longitudinal monitoring of adaptive behavior under pharmacologic pressure bridging the gap between molecular profiling and therapeutic response.

A major methodological advancement of this study is the development of a high-resolution, multiparametric 3D drug screening platform that integrates morphological, proliferative, and structural parameters to capture treatment responses beyond broadly used viability measurements. Conventional drug screening assays often rely on bulk luminescence or metabolic activity, masking sublethal phenotypes and adaptive cytostatic responses that preceded overt resistance. By contrast, our system, combining quantitative imaging with phenotype-integrated analytics, resolves the spatiotemporal heterogeneity of drug response within each PDO population. Using this two-tier, phenotype-integrated hit-calling framework, we found that KRAS mutational status emerged as a major determinant of therapeutic phenotype across our PDO cohort. KRAS wild-type PDOs predominantly underwent true cytotoxic collapse, characterized by structural disintegration and loss of volumetric integrity in which indicating increased vulnerability to single-agent therapies capable of inducing direct cellular disintegration. In contrast, KRAS-mutant PDOs predominantly exhibited cytostatic growth-arrest phenotypes without overt structural disintegration, reflecting a more tolerant drug response state. Clinically, this pattern closely mirrors the limited efficacy of single targeted agents in KRAS-mutant metastatic CRC and reinforces the rationale for combination-based therapeutic strategies^68^. These findings not only refine phenotypic drug-response profile in colorectal cancer but also highlight a need and the potential of multi-dimensional screening approaches as predictive tools for patient stratification.

Building on this functional landscape, to our knowledge, we present the first PDO-based characterization of resistance to the novel pan-KRAS inhibitor BI-2865, a next-generation compound that targets the GDP-bound inactive state of KRAS to block its activation across oncogenic variants^22^. Owing to its broad inhibitory spectrum and favorable pharmacologic profile, BI-2865 holds strong potential as a clinically translatable therapy for KRAS-driven cancers. Through long-term drug exposure, both KRAS^WT^ and KRAS^G13D^ PDO models developed resistance, albeit through distinct molecular and phenotypic trajectories. Resistance in both KRAS^WT^ and KRAS^G13D^ PDOs arose primarily through transcriptional plasticity rather than widespread genomic remodeling. The global mutational landscapes remained largely conserved, though selective enrichment of variants in *RNF12*, *DHPS*, *ZNF658*, and *BTNL3* suggested positive selection of subclones with enhanced stress tolerance, proteostatic control, and epithelial immune adaptability. These genetic shifts likely provided a permissive condition upon which widespread transcriptional reprogramming occurred. DNA barcoding analyses revealed that resistance in KRAS^WT^ organoids was predominantly driven by *de novo* emergent clones arising during drug exposure, reflecting transcriptional reprogramming and phenotypic adaptation, consistent with previous observations that drug-tolerant persister cells can evolve to acquire genetic resistance under sustained targeted inhibition^69^. In contrast, KRAS^G13D^ PDOs were dominated by pre-existing tolerant clones, indicative of Darwinian selection and expansion of pre-adapted subpopulations under therapeutic pressure, resembling the deterministic selection of pre-existing resistant subclones observed through large-scale barcoding in EGFR-mutant lung cancer models^70^. Transcriptomic profiling further revealed suppression of proliferative programs and activation of stress-response, cytoskeletal, and adhesion pathways characteristic of drug-tolerant persister states. The KRAS^WT^ organoids adopted a quiescent, adhesion-anchored phenotype stabilized by NF-κB–mediated signaling, whereas KRAS^G13D^ PDOs engaged Rho GTPase, WNT/β-catenin, PI3K/AKT–NF-κB, and compensatory ERBB/EGFR pathways, establishing a highly plastic, multi-layered resistance network. Functionally, the emergence of resistance in these models underscores the interchangeability of adaptive states and supports the concept that tumors navigate multidimensional fitness landscapes rather than linear genetic hierarchies.

Given the translational extension provided by functional drug screening, BI-2865 resistance reshaped the spectrum of actionable therapeutic vulnerabilities in a genotype-dependent manner. By specifically interrogating cytotoxic responses that emerge after resistance acquisition, this approach revealed dependencies that are unmasked by adaptation itself, underscoring resistance as an active evolutionary process rather than a passive loss of drug sensitivity. In KRAS^WT^ PDOs, resistance was associated with selective sensitivity to cell-cycle checkpoint and DNA damage response inhibitors, consistent with transcriptional suppression of proliferative programs, attenuated p53 signaling, and increased checkpoint dependence. The parallel acquisition of sensitivity to RTK–PI3K–mTOR inhibitors indicates that, despite growth arrest, KRAS^WT^ resistant cells retain a minimal requirement for residual survival signaling, supporting a quiescent, checkpoint-fragile persister-like state. By contrast, BI-2865-resistant KRAS^G13D^ organoids displayed a broader vulnerability landscape, reflecting a signal-amplified and highly plastic resistant state. Acquired sensitivities to RTK, SRC, MEK/ERK, and PI3K/AKT/mTOR inhibitors mirror transcriptional activation of these pathways, while the emergence of selective vulnerability to PORCN inhibition directly links WNT ligand–dependent regenerative signaling to a druggable resistance specific dependency.

Incorporation of single-cell and lineage-resolved analyses provides a mechanistic framework for understanding how these functional vulnerabilities arise. Rather than converging on a uniform resistant identity, KRAS^G13D^ PDOs maintain a shared transcriptional landscape within which multiple resistant trajectories coexist. Cluster-resolved pathway analysis reveals that resistance-associated signaling is organized in a modular and state dependent manner, with proliferative, metabolic, inflammatory, and stress-adaptive programs differentially deployed across transcriptionally defined clusters. Importantly, pathways such as MYC- and mTORC1-associated programs often represent stable, cluster-intrinsic states maintained under treatment, whereas plasticity-associated (WNT/TGF-β), inflammatory (TNFα/NFκB, interferon), and metabolic (oxidative phosphorylation, fatty acid metabolism) modules are selectively enriched in distinct resistant clusters. As a result, the same pathway may be upregulated in one resistant state while being attenuated in another, reflecting functional partitioning rather than conflicting signaling behavior. Single-cell pathway scoring further demonstrates that these programs are broadly distributed across clonal classes, indicating that resistance is sustained through redistribution of pre-existing cellular states rather than dominance of a single clonal lineage. Together, this state-resolved architecture resolves the molecular heterogeneity observed at the single-cell level with the emergence of genotype-specific therapeutic vulnerabilities, highlighting that effective strategies to overcome pan-KRAS inhibitor resistance will likely require combination therapies tailored not only to oncogenic genotype, but also to the spectrum of resistant cell states present within a tumor. In summary, our data support a model in which resistance to pan-KRAS inhibition is not defined by a single dominant signaling pathway or clonal solution, but instead emerges through state-resolved redistribution of pre-existing cellular programs. This modular resistance architecture enables parallel adaptive strategies, including checkpoint-dependent stress tolerance, epithelial plasticity, inflammatory signaling, and metabolic reinforcement, to coexist within a shared transcriptional landscape. By linking these state-specific programs to genotype-dependent therapeutic vulnerabilities, our findings highlight the importance of integrating functional drug screening with single-cell–resolved molecular profiling to rationally design combination strategies capable of targeting heterogeneous resistant populations.

Despite these advances, number of limitations should be acknowledged. while PDOs robustly model tumor-intrinsic responses, they do not capture extrinsic modulators such as immune surveillance, stromal signaling, or pharmacokinetic constraints that can influence therapeutic outcomes in vivo. Future studies incorporating PDO–immune co-cultures, organ-on-chip vascular interfaces, and longitudinal sampling during clinical treatment could address these gaps and validate the predictive potential of PDO-derived phenotypic biomarkers. Additionally, integrating single-cell chromatin accessibility (scATAC-seq) and spatial transcriptomics could delineate how epigenetic remodeling contributes to the persistence of tolerant states during drug holidays or sequential therapy. Nonetheless, the present study demonstrates that combining PDO-based experimentation with multi-parametric analytics and clonal tracing provides a mechanistically rich, scalable, and patient-relevant platform for understanding the evolutionary basis of drug resistance. Such integrative frameworks will be key for the rational design of next-generation combination therapies capable of converting adaptive plasticity from a dependency into a therapeutic vulnerability.

## Materials and Methods

### Ethical regulations

PDOs from metastatic colorectal cancer were established using fresh tumor specimens collected primarily from patients with colorectal cancer who had developed liver metastases and underwent surgical resection at either Hacettepe University Hospital or Ankara City Hospital. Ethical approval for the study was granted by the Hacettepe University Non-Interventional Clinical Research Ethics Committee (decision number: KA-20136).

### Establishing patient-derived metastatic organoids

The establishment and maintenance of metastatic colorectal cancer organoids were carried out with modifications based on previously published protocols, including those described by Sato et al. (2011), van de Wetering et al. (2015), and Vlachogiannis et al. (2018), which outline methodologies for the derivation of organoids from primary and metastatic colorectal tumors^7,67,71^. Following surgical resection, tumor specimens were promptly immersed in ice-cold Dulbecco’s Phosphate Buffered Saline (DPBS; Sartorius) containing 1× Penicillin-Streptomycin (Thermo Fisher Scientific) and transported to the laboratory. The tissue was rinsed three times with chilled DPBS containing 1× Penicillin-Streptomycin (Thermo Fisher Scientific) and Primocin 100 μg/ml (InvivoGen), to remove blood and debris, then mechanically minced into small fragments with bisturi. Subsequently, the fragments were incubated in DPBS supplemented with 5 mM EDTA (Sartorius) for 15 minutes at room temperature to facilitate cell dissociation. Enzymatic digestion was then performed by incubating the tissue fragments with 1X TrypLE Select Enzyme (Thermo Fisher Scientific) at 37 °C for 60 minutes. Following digestion, gentle pipetting was applied to enhance dissociation. The resulting cell suspension was filtered through a 40 µm cell strainer (Greiner) to remove residual undigested material and centrifuged at 1200 rpm for 5 minutes at 4 °C. The pelleted cells were resuspended in 120 µl of ice-cold growth factor–reduced Matrigel (Corning, #356231) and plated in 24 well plate.

The Matrigel domes were allowed to solidify in a humidified incubator at 37 °C with 5% CO₂ for 20 minutes before being overlaid with 500 µl of organoid culture medium. The basal medium consisted of Advanced DMEM/F12 (Thermo Fisher Scientific) supplemented with 1× Penicillin-Streptomycin, 2 mM L-Glutamine (Thermo Fisher Scientific), 100 μg/ml Primocin (InvivoGen), and the following additives and growth factors: 1× B27 and 1× N2 (Gibco), 0.01% BSA (Sigma), 4 mM Nicotinamide (Sigma), 10 nM Gastrin (Sigma Aldrich), 50 ng/ml human EGF, 100 ng/ml human Noggin, 500 ng/ml human R-Spondin-1, 10 ng/ml human FGF-10, 10 ng/ml FGF-basic (all from BioLegend), 0.5 µM A83-01 (Tocris), 100 ng/ml recombinant Wnt-3a (R&D Systems), 0.5 µM A83-01 (Tocris), 1 µM Prostaglandin E₂ (Tocris), and 5 µM SB202190 (Sigma Aldrich). For the initial establishment phase, the ROCK inhibitor Y-27632 (10 µM; Sigma Aldrich) was included to promote cell survival and attachment. Medium was replaced every 48 hours, and the cultures were monitored microscopically for organoid formation from single cells or cell aggregates.

### Histopathology and immunohistochemistry

Histological and immunohistochemical characterization was performed on both PDOs and matched primary tumor tissues. To preserve their 3D architecture, PDOs were recovered from Matrigel using ice-cold Cell Recovery Solution (Corning) for 1 hour at 4°C, washed in cold PBS, centrifuged (4000 rpm, 4 min, 4°C), and fixed in 4% paraformaldehyde (Sigma Aldrich) for 1 hour. The fixed organoids were embedded in 2% low-melting agarose, followed by paraffin embedding. Sections (3-4 μm) were stained with hematoxylin and eosin (H&E) and examined by light microscopy. For immunohistochemistry, additional paraffin sections were prepared from PDOs. After standard deparaffinization and antigen retrieval, staining was performed using antibodies against Ki-67 (proliferation), CK20 (epithelial marker), and CDX-2 (intestine-specific transcription factor). Visualization was achieved using HRP-conjugated secondary antibodies and DAB chromogen.

### WES

Genomic DNA was extracted from primary tumor tissues and matched peripheral blood samples using the PureLink Genomic DNA Mini Kit (Invitrogen), and from organoid cultures using the Quick-DNA Microprep Kit (Zymo Research), following the manufacturers’ protocols. Whole-exome libraries were prepared using the Twist Comprehensive Exome kit and sequenced on the Illumina NovaSeq 6000 platformSomatic single-nucleotide variants (SNVs) and small insertions/deletions (indels) were identified using the nf-core/sarek v3.4.0 pipeline (ref1). The workflow included initial quality assessment using FastQC v.0.12.1 followed by aggregation with MultiQC v1.17. Because sequencing reads showed consistently high-quality metrics across all samples (per-base quality > Q30 and detected adapter contamination < %1), no trimming step was applied, and reads were directly aligned to the GRCh38 human reference genome using BWA-MEM v.0.7.17.

Resulting variant call format (VCF) files were annotated with snpEff (GRCh38.105 database). Annotated VCFs were imported into R (v4.4.0) using vcfR v1.15.0 for downstream filtering and visualization. Only PASS, exonic, and non-synonymous variants (missense, nonsense, frameshift, in-frame indels, and splice-site variants) were retained for analysis. Gene-level summaries and heatmaps for each tumor–PDO pairs were generated using ComplexHeatmap v2.20.0.

To evaluate whether specific somatic variants were enriched or depleted during the acquisition of BI2865 resistance, we compared variant allele frequencies (VAFs) between resistant PDOs (PDO-01 and PDO-06) and their matched DMSO controls. For each SNV, reference and alternate read counts were aggregated per replicate, and variants with insufficient sequencing depth (<5 total reads per sample) were excluded. Statistical enrichment was tested per variant using a stratified framework: when two or more resistant replicates had a measurable coverage, Cochran–Mantel–Haenszel (CMH) test was applied to compare ALT/REF counts across replicates while controlling for replicate-specific variation; otherwise, Fisher’s exact test was used. Resulting p-values were adjusted (Benjamini–Hochberg), and variants with FDR < 0.05 were retained as significant. Variants were classified as treatment-enriched if they showed (i) padj < 0.05, (ii) an increase in allele frequency of at least ΔVAF ≥ 0.2 in all three replicates. Conversely, variants with padj < 0.05 and ΔVAF ≤ –0.2 were considered DMSO-enriched (lost in resistance). Variants were visualized using DMSO-versus-treatment VAF scatterplots.

Copy-number alterations (CNAs) were derived using CNVkit, as implemented in the nf-core/sarek pipeline. For each sample, CNVkit was run in WES mode. A pooled reference was constructed from the matched normal samples by the Sarek workflow unless otherwise specified. CNVkit produced both bin-level coverage files (.cnr) and segmented copy-number files (.cns) for each tumor and PDO sample.

To correct copy-number estimates for tumor purity and ploidy, we applied FACETS v0.16 on recalibrated BAM files for each tumor–normal pair. FACETS outputs provided estimates of sample purity, total ploidy, and allelic imbalance. These purity values were subsequently used to adjust CNVkit segment-level estimates through CNVkit’s call function, enabling generation of purity-normalized cns files.

Gene-level copy-number calls were derived using CNVkit by integrating purity-adjusted cns files with bin-level cnr data. To focus on high-confidence copy-number alterations in cancer-relevant genes, we applied additional filtering at the gene level on the CNVkit genemetrics output. For each tumor and PDO sample, genes were retained only if they showed an absolute log₂ copy-number ratio ≥ 1.2, were supported by >10 probes and >500 segment_probes, and had either a mean depth ≥ 100 or a CNVkit weight ≥ 30. Segments whose confidence intervals crossed zero (ci_lo > 0 and ci_hi > 0) were excluded to remove uncertain events. From these filtered calls, we extracted genes belonging to a curated panel of colorectal cancer– and cancer-related genes (including APC, KRAS, TP53, SMAD4, EGFR, PIK3CA, and others; full list in Table S[X]), and used these high-confidence gains and losses are visualized using ComplexHeatmap together with SNVs.

Genome-wide copy-number profiles were visualized by converting CNVkit .cnr files into a unified matrix of bin-level log₂ copy-number ratios. For each sample, the .cnr files were imported and merged into a long-format table containing the genomic bin coordinates and corresponding log₂ values. Log₂ ratios were then limited within the range −1.2 to +1.2 to prevent extreme values from dominating the color scale. This matrix was plotted using the ComplexHeatmap package, where each row represents a sample and each column represents a genomic bin, producing a genome-wide heatmap of copy-number gains (red) and losses (blue) for tumor and PDO samples.

### Bulk-RNA Seq

Total RNA was isolated from patient-derived organoids and matched primary tumor tissues using the Zymo Quick-RNA Microprep Kit and Qiagen RNeasy Mini Kit, respectively. RNA quality was assessed using RIN values. Libraries were prepared using the Illumina Stranded mRNA Prep Kit with poly-A selection, and sequencing was performed on the Illumina NovaSeq 6000 platform to generate 150 bp paired-end reads.

Gene count matrices for each sample were obtained by implementing the nf-core/rnaseq v.3.18.0 pipeline (ref2). FastQC v.0.12.1 and Trim Galore v.4.9 was used to trim sequencing reads, eliminating the remains of Illumina adaptors and discarding reads that were shorter than 20 bp. Then, reads were aligned to the GRCh38 reference genome using STAR v.2.7.10a together with the corresponding Ensembl transcript annotation. Gene-level expression was quantified using RSEM v1.3.1, which generated estimated gene counts as well as transcript-per-million (TPM) values. Downstream analysis of raw merged gene counts was carried out in R.

Since tumor and organoid libraries were generated in two separate sequencing batches, batch effects were corrected using ComBat-seq function of sva package v.3.52.0 by specifying the batch variable corresponding to “tumor” and “organoid” sample groups. The batch-corrected counts were transformed using variance-stabilizing transformation (VST) from DESeq2 v.1.44.0. Mean expression for each gene was computed as the average VST value across all samples, and genes with a mean expression below 1 were excluded to remove low-abundance, noise-prone features. Gene-level variability was quantified using the coefficient of variation (CV), calculated as the standard deviation divided by the mean expression across samples. The 8,000 most variable genes were selected based on CV and for each matched tumor–PDO pair, genes with elevated expression (z-score > 1) in both samples were identified to examine shared transcriptional features. Results were visualized as a heatmap using Pheatmap v.1.013.

Dimensionality reduction analyses were performed using UMAP on the z-score–scaled expression matrix to evaluate global transcriptional similarity between tumor–PDO pairs, and pairwise similarity was further quantified by computing Pearson correlation coefficients from the same scaled data.

Differential gene expression analysis of BI2865-resistant PDOs (PDO-01 and PDO-06) was performed separately against their matched DMSO controls. RSEM gene-level counts were filtered to remove genes with fewer than 10 counts in at least 50% of samples. DESeq2 was used to construct a count-based model with treatment (control vs. resistant) as the design variable, and differential expression testing was performed using the Wald test. Genes with an adjusted p-value (padj < 0.01) were considered significantly up- or down-regulated. Variance-stabilizing transformation (VST) was applied for visualization and PCA.

Significantly enriched pathways were defined using an FDR cutoff of 0.05 with clusterProfiler v.4.12.6. Enriched biological processes and signaling pathways were visualized using dot plots. Pathway-level alterations were evaluated using GSVA v.1.52.3 with MSigDB v.24.1.0 C6 (oncogenic signatures) gene sets. GSVA enrichment scores were contrasted between resistant and control samples using limma v.3.60.6 with empirical Bayes moderation.

### 3D-image-based drug screening

PDOs were embedded in 10 µl of growth factor–reduced Matrigel (Corning) per well in PhenoPlate 384-well black 6057302 (Revvity) to ensure optimal three-dimensional support and compatibility with high-content imaging. A seeding density of 1,500 cells per well was determined as optimal for maintaining organoid viability and growth over a 10-day assay period, based on morphological assessment and response consistency to the positive control drug, Nocodazole.

Following cell seeding, culture medium was replaced on day 2, and treatment compounds were administered on days 4 and 7 using a library of 50 clinically relevant compounds either in use or under investigation in phase I–III clinical trials was screened at a final concentration of 1 µM. Prior to imaging, organoids were incubated with 1 µM EdU (5-ethynyl-2’-deoxyuridine) for 4 hours to label cells that undergo effective DNA replication during the labeling window, followed by fixation in 4% paraformaldehyde for 20 minutes at room temperature. Subsequently, organoids were stained with SYTOX™ Green Nucleic Acid StainS7020 (Life technologies) for nuclear visualization HCS cell mask deep red H32721 (Life technologies) for volumetric staining, and Click-iT EdU Cell Proliferation Kit for Alexa Fluor 594 C10339 (Life Technologies) for detection of EdU incorporation. High-content 3D imaging was performed using the Operetta CLS confocal imaging system (Revvity), equipped with a 20× water-immersion objective (NA = 1.0). From each well, five representative fields were imaged, and Z-stack acquisition was performed at 1 µm intervals to allow full-volume 3D reconstruction of individual organoids. Image analysis was performed using Harmony (v4.9, PerkinElmer) software. For each organoid, nuclear count (cellularity), total volume, and the number of EdU-positive nuclei were quantified to evaluate drug-induced effects on proliferation and viability Quantitative results from Harmony were exported at the single-organoid (object) level, enabling multiparametric analysis of individual organoids. Object-level measurements were then summarized across technical replicates for each drug–PDO–condition combination. For each result, values were normalized to the corresponding DMSO control wells on the same plate to yield percent-of-control responses. Normalized volume, nuclei and EdU measurements for 10 PDOs (PDO-01, PDO-02, PDO-03, PDO-04, PDO-06, PDO-13, PDO-15, PDO-16, PDO-17, PDO-20) under DMSO and BI-2865 treatment were assembled into a drug (rows) × PDO (columns) matrix and visualized as heatmaps, with color intensity reflecting organoid volume relative to control. Drugs were annotated with their primary mechanism of action and grouped accordingly in the heatmap. In addition, drugs were classified as cytotoxic when either all three measurements (volume, nuclei, and EdU) or two measurements (volume and nuclei) were reduced to <50% of DMSO control, and as cytostatic when only the EdU-positive fraction was reduced to <50% of control while volume and nuclear count remained ≥50%. This classification was indicated by colored markers alongside each PDO column.

### Barcoding PDOs

Clonal barcoding of individual organoid cells was performed using the CloneTracker XP™ 1M Barcode-3’ Library Pool 1 with RFP-Puro selection cassette (Cellecta, USA; Lot# 180817002). This lentiviral-based library enables heritable labeling of single cells through stable genomic integration and transcription of unique barcode sequences, thereby allowing longitudinal clonal tracking and transcriptomic profiling.

To ensure single-barcode integration per cell, lentiviral transduction was performed at a multiplicity of infection (MOI) of 0.1 in the presence of 0.8 µg/ml polybrene, corresponding to an estimated transduction efficiency of ∼10%. A total of 1 × 10⁶ cells derived from the Pat-1 and Pat-6 PDOs were transduced according to the manufacturer’s protocol. After overnight incubation in suspension with virus-containing medium, cells were pelleted, resuspended in 500 µl of growth factor–reduced Matrigel (Corning), and seeded into five wells of a 24-well plate (2 × 10⁵ cells per well in 100 µl Matrigel overlaid with 500 µl medium).

Selection was initiated with puromycin (1.5 µg/ml), a concentration determined via prior kill curve analysis, and continued until all non-transduced control cells were eliminated. Successfully barcoded organoid populations were expanded under standard culture conditions until reaching sufficient numbers for downstream experimental applications.

### Generation of pan-KRAS inhibitor-resistant organoids

To model the development of acquired resistance, two PDOs, a KRAS wild-type (PDO-01) and a KRAS G13D mutant (PDO-06) were treated to continuous drug exposure with the novel pan-KRAS inhibitor BI-2865 (MedChem). For each PDO, 1 × 10⁶ cells were seeded in 12-well plates (three biological replicates per line), embedded in 500 µl growth factor–reduced Matrigel (Corning) and overlaid with 1 ml of complete organoid culture medium. Organoids were allowed to expand for 4 days, after which the culture medium was replaced with fresh medium supplemented with BI-2865.

Drug treatment was initiated at a concentration 10-fold below the previously determined IC₅₀ value for each line. With every subsequent medium change, the BI-2865 concentration gradually increased in a stepwise manner, doubling the dose each time to mimic escalating therapeutic pressure. This adaptation strategy was continued over a period of approximately 3 months for both PDO lines. DMSO was used in parallel cultures as a vehicle control throughout the selection period. During this time, organoids were monitored for viability, morphology, and recovery kinetics following each dose escalation to ensure gradual enrichment of drug-tolerant subclones.

### Barcode amplification and next generation library preparation

Following the development of acquired drug resistance, genomic DNA was extracted from organoid cultures at the experimental endpoint using the Zymo Quick-DNA Microprep Kit, according to the manufacturer’s instructions. For barcode sequencing, the integrated lentiviral barcode regions were PCR-amplified following the standard protocol provided by Cellecta^TM^. The amplified barcode libraries, containing variable sequences of 14 to 30 bp in length, were prepared for next-generation sequencing. Paired-end 150 bp sequencing was performed on the Illumina MiSeq platform to ensure accurate capture of the barcode diversity.

### Barcode bioinformatics analysis

Barcode regions (∼14–30 bp variable sequences) were amplified and sequenced using 150 bp paired-end reads on the Illumina MiSeq platform. Sequencing libraries included samples from both BI-2865-resistant PDO populations and matched DMSO-treated controls. Raw FASTQ files were subjected to quality control using FastQC, and reads with a Phred quality score below 20 were excluded from downstream analysis. Adapter and constant flanking regions were trimmed using Trimmomatic v.0.39, resulting in a 48 bp sequence corresponding to the unique barcode.

A synthetic reference library of 1 million possible barcodes was generated in FASTA format using the R package insect, based on the known design of the Cellecta CloneTracker XP system. This reference was indexed with Salmon v1.9.0k-mer size of 27. Then, barcode-specific transcript quantification was performed using Salmon, and output read counts were exported for downstream analysis.

Barcode growth dynamics were calculated by comparing barcode abundances between drug-treated and control samples. Barcodes undetected in the DMSO sample but present in resistant replicates were assigned a nominal count of 1 in the DMSO dataset to allow calculation of frequency ratios. For each barcode, growth rates were calculated by dividing its frequency in each resistant replicate by its corresponding frequency in the DMSO control. Based on these growth factors, barcodes were functionally classified into three different categories: Pre-existing: barcodes showing positive growth and detected in ≥2 resistant replicates; De novo: barcodes with positive growth detected in only a single resistant replicate; Sensitive: barcodes whose abundance decreased relative to DMSO following drug treatment. This classification enabled resolution of clonal selection dynamics and identification of adaptation patterns under panKRAS inhibitor pressure. Each barcode was given a unique color, and sample-wise frequency distributions were displayed as bar plots.

### Dose response curves

To assess the development of resistance to BI-2865 in PDO-06 organoids, 3D cell viability assays were performed. DMSO-treated controls, including both the initial population and a non-barcoded control, together with three independent BI-2865-resistant replicates (ResA, ResB, and ResC), previously cultured in 12-well plates, were harvested and enzymatically dissociated into single cells using TrypLE Express (Thermo Fisher Scientific). A total of 1.5 × 10³ cells per well were embedded in 10 μl of growth factor–reduced Matrigel and seeded into 384-well plates, overlaid with 50 μl of complete organoid medium, and included in dose–response viability curves.

Organoids were allowed to re-form and expand for four days, with a medium refresh at 24 hours post-seeding. Subsequently, BI-2865 was administered in a range of concentrations and maintained for six days to generate dose–response curves. The medium was replaced once after 72 hours. At the end of the treatment period, cell viability was quantified using the CellTiter-Glo® 3D Cell Viability Assay (Promega), and half-maximal inhibitory concentrations (IC₅₀) were calculated for each condition.

Dose–response curves were generated by normalizing raw viability measurements within each condition to the maximal viability observed across all doses and scaling them to percent viability relative to DMSO controls. For each concentration and condition, technical replicates were averaged, and the standard error of the mean was calculated. Dose–response relationships were then modeled using a five-parameter log-logistic (LL.5) function implemented in the drc (v3.0-1) R package, enabling estimation of smooth sigmoidal curves for all conditions. Plots display mean normalized viability with error bars together with the corresponding fitted dose–response curves for each condition.

### Single-cell RNA Profiling

Single-cell RNA sequencing (scRNA-seq) was performed on PDO samples using the DNBelab C4 Series Single-Cell Library Prep Set (MGI, #1000021082), following the manufacturer’s standard protocol. Organoids were enzymatically dissociated into single cells using 1X TrypLE (Thermo Fisher Scientific) at 37 °C, followed by centrifugation. After centrifugation, the cell pellet was resuspended in PBS containing 0.04% BSA and passed through a 35-µm cell strainer (Corning, #352235) to obtain a high-quality single-cell suspension.

Cells and barcoded magnetic beads were prepared at a final concentration of 1000 cells/μL and 1000 beads/μL, respectively. For each sample, 100,000 single cells and an equal number of barcoded beads were loaded into the DNBelab C4 microfluidic chip for droplet encapsulation. After on-chip droplet formation and incubation at room temperature to allow for mRNA capture, droplets were broken, and beads were collected for reverse transcription and subsequent cDNA amplification. Final libraries were prepared according to MGI protocols and subjected to high-throughput paired-end sequencing (150 bp) on the MGISEQ-T7 platform (MGI Tech, China).

Raw fastq files from DMSO-control and BI2865-resistant PDO-06 ResC samples were processed using the DNBelab C Series HT scRNA-analysis pipeline (https://github.com/MGI-tech-bioinformatics/DNBelab_C_Series_HT_scRNA-analysis-software). Reads were quality-filtered, barcodes and UMIs were parsed, and reads with Phred <20 or mapping quality <20 were removed. High-quality reads were aligned to GRCh38 using STAR, and processed with Sambamba and PISA for sorting, filtering, and generating feature counts. UMI correction was performed using Hamming-distance collapse to retain the highest-frequency UMI. Gene annotations were applied to produce a raw gene–cell matrix.

Oligo barcodes underwent QC and were merged with cDNA barcodes via similarity matching. Cell calling was used to identify true cells and remove background and low-quality barcodes. The resulting filtered expression matrix was carried forward for clustering and downstream analyses.

Downstream filtering and quality control steps were carried out in R. Ambient RNA contamination was removed using SoupX v.1.6.2, which estimated background RNA levels from raw and filtered matrices and generated corrected count matrices for each sample. The corrected datasets were then processed with DoubletFinder v.2.4.0 to identify and exclude predicted doublets. Only high-confidence singlet cells from each condition were retained and merged into a unified Seurat object for downstream analysis. Low-quality cells were removed by retaining only cells with >1,000 detected genes and <25% mitochondrial transcript content. Quality control metrics before and after filtering are provided **(Supplementary Fig. 4).**

Data processing and dimensionality reduction were performed using Seurat v5.3.0. Count matrices were normalized, and highly variable genes were identified for downstream analysis. Gene expression values were scaled, and a principal component analysis was performed to capture major sources of variation. The first 30 principal components were used to construct a nearest-neighbor graph and generate a two-dimensional UMAP embedding for visualization of cellular heterogeneity. Cells were assigned cell-cycle scores based on canonical S phase and G2/M phase gene sets, allowing estimation of their relative stage in the cell cycle. These cell-cycle states were then visualized for each cluster and compared between DMSO and ResC groups to assess potential treatment-related shifts. To statistically evaluate whether the distribution of G1, S, and G2/M phases differed between conditions within each cluster, a chi-square test was applied.

### Detection of Expressed Barcodes in the Single Cell Data

Expressed barcode quantification from single-cell RNA-seq data was performed using the *simpleaf* implementation of the Salmon–Alevin-fry workflow (simpleaf v0.19.5, alevin-fry v0.11.2) together with Salmon-based indexing. The same Salmon index generated for DNA barcode calling was reused for scRNA-based barcode assignment. A custom chemistry definition was added to simpleaf to accurately represent the library structure, specifying that read 1 contains a 20-bp cell barcode followed by a 10-bp UMI (geometry: 1{b[20]u[10]x:}2{r:}), while read 2 contains the cDNA sequence.Raw paired-end FASTQ files were processed in *quant* mode using this custom chemistry, the barcode index, and an explicit whitelist of high-confidence cell barcodes. Cell barcodes identified by the DNBelab C Series HT pipeline were exported as barcode count tables; the count column was removed, and the resulting list of barcodes was supplied to simpleaf/alevin-fry via the --explicit-pl option to ensure that only biologically valid barcodes were retained during quantification. A transcript-to-gene mapping file (tgMap) was generated in which each synthetic barcode transcript (IDs 1–1,000,000) was mapped to itself, ensuring that Alevin-fry treated each barcode as an independent gene during quantification. Barcode assignment was performed using the *cr-like-em* resolution model.

Following alevin-fry quantification, count matrices were imported into R using fishpond v2.10.0 and converted into Seurat objects. To determine the expressed barcode for each cell, barcode-level UMI counts were examined and the dominant barcode was selected based on several quantitative thresholds. First, a barcode was considered valid only if it accumulated at least 2 UMIs (minUMI), ensuring that single-UMI noise was not misinterpreted as true expression. Second, the top barcode was required to contribute at least 70% of all barcode-derived UMIs in that cell (fractional dominance; minFrac = 0.7), ensuring that the assigned barcode clearly outweighed all others. Third, the ratio between the top barcode and the second-highest barcode had to be at least 2-fold (minRatio = 2), preventing ambiguous assignments when two barcodes were expressed at similar levels.

Cells in which more than three different barcodes were detected (maxDetected = 3) were labeled as multiplets, unless the top barcode showed extreme dominance—defined as a ≥10-fold excess over the second barcode (ratio ≥10) while having ≤50 detectable barcodes—allowing confident recovery of strong single-barcode signals even in noisy cells. Cells failing any of these criteria were classified as ambiguous or unassigned.

Barcode translation tables generated from the DNBelab pipeline were then used to link each expressed synthetic barcode to its corresponding individual cells. The results were incorporated into the metadata of the merged Seurat object containing both DMSO and BI-2865–resistant (ResC) samples, enabling downstream lineage-tracking and transcriptional analyses. Subsequent analyses were restricted to cells with a successfully assigned synthetic barcode, and cells lacking an assigned barcode were excluded.

Expressed barcode frequencies from single-cell data were quantified by aggregating per-cell barcode assignments across conditions. For each sample, cells with a valid expressed barcode were counted per barcode, and barcode frequencies were calculated by normalizing counts to the total number of barcode-assigned cells within that sample. DNA-derived barcode classifications obtained previously for the BI-2865–resistant population (pre-existing, de novo, sensitive) were mapped onto the corresponding expressed barcodes detected in single-cell RNA-seq. Barcodes detected only in the single-cell dataset but absent from DNA profiling were labeled as “single-cell–specific”. The same color scheme used in DNA-barcode visualizations was applied to ensure consistency between datasets. Frequencies of classified barcodes were plotted for DMSO control and ResC samples, enabling comparison of barcode frequencies at the single-cell transcriptional level.

UMAPs of barcode-labeled cells from BI-2865–resistant (ResC) and DMSO samples are shown split by transcriptional cluster, with cells colored according to their assigned barcode class (de novo, pre-existing, sensitive, or single-cell specific). This visualization highlights the spatial distribution of lineage classes across clusters and conditions. Corresponding stacked barplots depict, for each cluster, the fraction of cells belonging to each barcode class in DMSO and ResC samples, enabling quantification of shifts in barcode-class composition between conditions at the cluster level.

Expression of intestinal lineage marker genes was visualized using a dot plot in Seurat. For each cluster–condition group, Seurat’s DotPlot function was used to display the average normalized expression of each marker (color scale) and the percentage of cells expressing that marker (dot size).

### Differential Gene Expression (DEG) Analysis

In order to reveal distinct resistance mechanisms across transcriptional clusters and conditions, cluster-level differential expression and Hallmark pathway enrichment analyses were performed. Differential expression was computed separately for each cluster–condition identity (cluster_sample: clusters 0–4 in DMSO vs ResC), requiring genes to be expressed in at least 25% of cells and to exhibit an absolute log₂ fold change > 0.25. Genes with an adjusted p-value < 0.05 were retained and separated into upregulated and downregulated sets for each cluster. Hallmark pathway enrichment was then performed on these cluster-specific gene sets using clusterProfiler’s compareCluster function with MSigDB Hallmark gene sets (H collection, via msigdbr) as the reference. Enrichment results were visualized as dot plots for upregulated and downregulated genes separately, where dot color represents adjusted p-value and dot size reflects gene ratio, thereby highlighting pathway programs associated with resistance across clusters and between DMSO and ResC samples.

Per-cell pathway activity scores were computed for selected Hallmark gene sets (e.g. TNFα signaling via NFκB, MYC targets V1, MTORC1 signaling and fatty acid metabolism). For each pathway, the corresponding member genes were extracted from the enrichment results and a module score was calculated for every cell as the average expression of pathway genes relative to a background gene set using AddModuleScore function of seurat. Violin plots displaying the distribution of module scores per cluster_sample, split by barcode class, were used to visualize how pathway activity varies between barcode classes and treatment conditions at single-cell resolution.

## Supplementary Table and Figure Legends

**Supplementary Table 1. SNVs in tumor samples and matched PDOs. Supplementary Table 2. Clinical information of the patient cohort**

**Supplementary Figure 1. Chromosome-level copy number variation (CNV) profiles of tumors and matched patient-derived organoids (PDOs).**

**Supplementary Figure 2. Optimization of puromycin selection and lentiviral barcode transduction conditions in PDOs.**

**Supplementary Figure 3. Cell-cycle composition of BI-2865–resistant clusters compared with DMSO controls in PDO-06.**

**Supplementary Figure 4. Quality control metrics before and after filtering of single-cell RNA sequencing data.**

## Acknowledgments

Ahmet Acar was supported by the International Fellowship for Outstanding Researchers Program administrated by The Scientific and Technological Research Council of Türkiye (TÜBİTAK) (Grant number: 118C197). AA would also like to acknowledge Turkish Academy of Sciences Young Investigators Program (TÜBA-GEBİP), Science Academy Young Scientists Award Program (BAGEP), and Republic of Türkiye The Council of Higher Education Research Universities Support Program (Grant number: ADEP-108-2022-11202). Gizem Damla Yalcin was supported by the TÜBİTAK Directorate of Science Fellowships and Grant Program (BİDEB) under the 2211-C Program. We thank Mr. Ziya Birinci for sharing his expertise in histopathological analyses and for his valuable contributions to the staining experiments. We thank Roman Yunes from MGI Tech, Omer Comez from Invitrotek, and Bora Ergin from Intergen for their contributions to the single-cell RNA sequencing studies. We would like to thank Ms. Irem Bayram for her design of figures and past and present members of the Acar Lab for their valuable discussions and insights throughout the project.

## Data Availability

The data that supports the findings of this study are available from the corresponding author upon reasonable request.

## Author’s contributions

G.D.Y. designed experiments, performed data generation and interpreted the results. K.C.Y. designed bioinformatics pipelines for genomic data analysis and interpreted the results. R. K. and L. B. supported and supervised drug screening experiments. O. K., D. O. T, and C. S. performed pathological evaluation and provided support on histopathological examination. O. H. A., A. B. D., V. O., E. B. B., and S. Y. contributed to fresh tumor sample collection and the clinical interpretation of the results. A. A. conceived, designed, and supervised the study. A. A., G.D.Y., and K.C.Y. wrote the manuscript. All other authors contributed to the manuscript writing.

