## Supplementary File for "Molecular Profiling and High-Content Drug Screening of Metastatic Colorectal Cancer Organoids Reveal Evolutionary Mechanisms of Pan-KRAS Inhibitor Resistance"

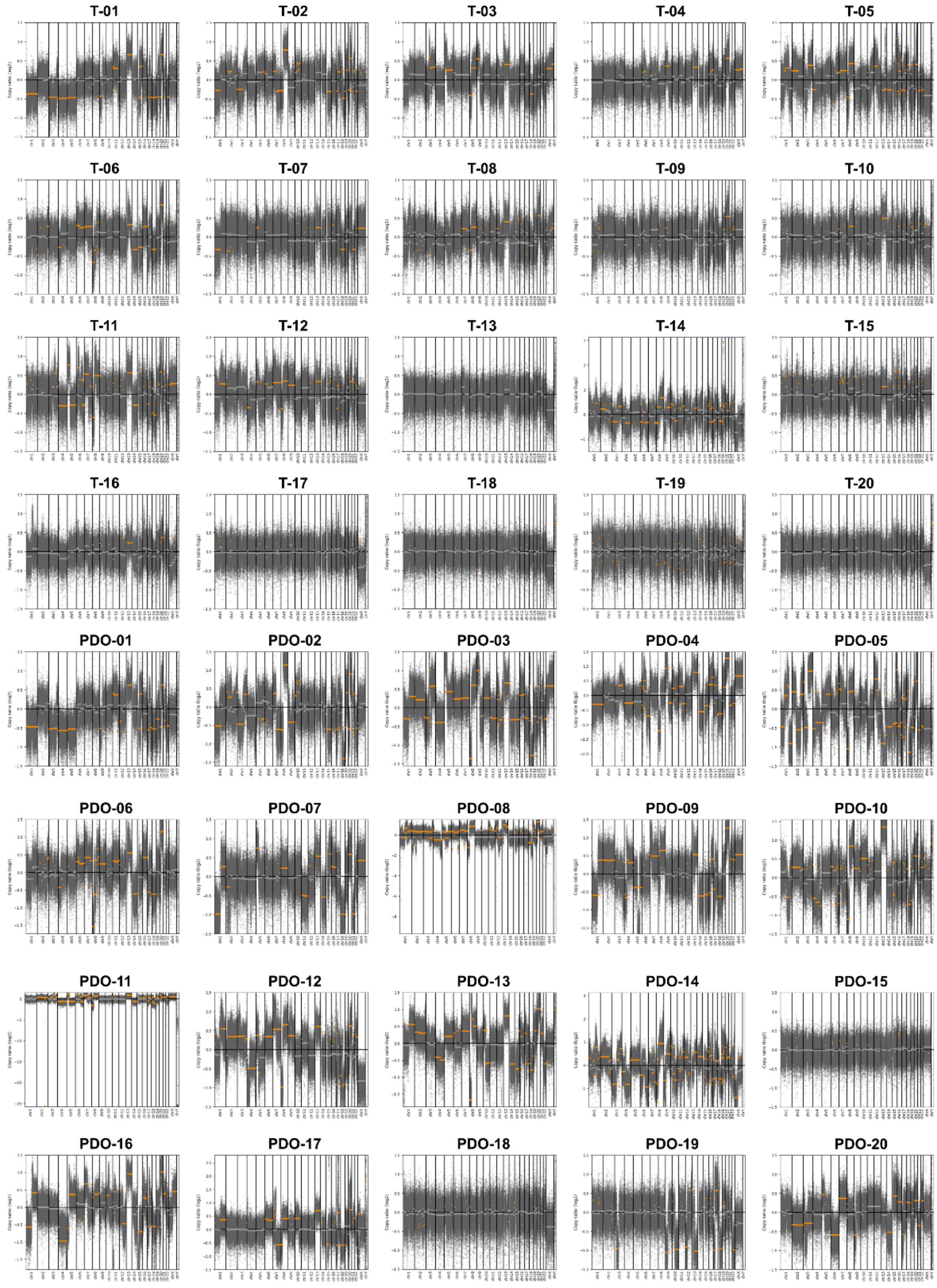

**Supplementary Fig 1. Chromosome-level copy number variation (CNV) profiles of tumors and matched patient-derived organoids (PDOs).** Genome-wide copy number alterations for all 20 primary tumors and their corresponding PDOs are shown as log2 copy number ratios plotted across all chromosomes (chr1–chrY). Each dot represents a genomic segment, with grey points indicating genome-wide CNV signals and orange points highlighting significantly altered regions. Chromosomal boundaries are indicated by vertical black lines. The horizontal dashed line denotes the diploid baseline (log2 ratio = 0), with positive and negative deviations corresponding to copy number gains and losses.

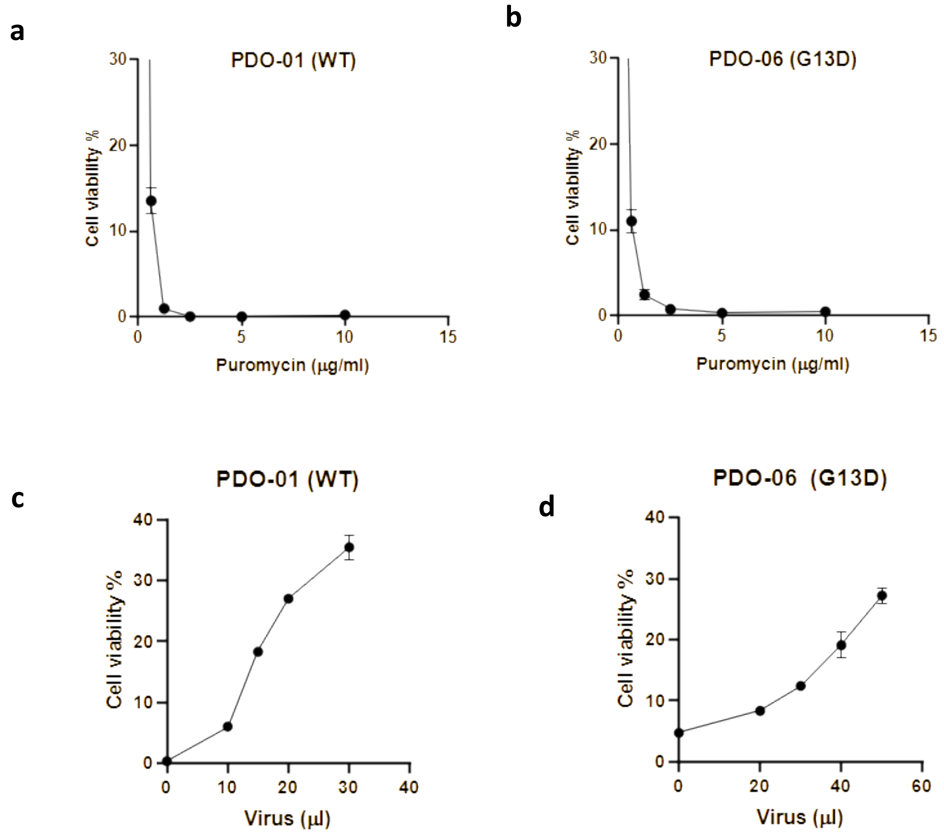


**Supplementary Fig 2. Optimization of puromycin selection and lentiviral barcode transduction conditions in PDOs. (a–b)** Puromycin sensitivity assays were performed in PDO-01 (a) and PDO-06 (b) to establish effective selection conditions. Following lentiviral induction, organoids were exposed to puromycin for 6 days, and cell viability was measured across increasing antibiotic concentrations. These analyses were used to define the minimal puromycin dose required for efficient elimination of non-transduced cells, which was determined as 0.5 µg/mL for PDO-01 and 1 µg/mL for PDO-06. **(c–d)** Optimization of lentiviral barcoding conditions were conducted in PDO-01 (c) and PDO-06 (d) by titrating the volume of lentiviral supernatant. Organoid viability following puromycin selection was quantified to estimate the viral input achieving efficient transduction while limiting multiple barcode integrations per cell. All experiments were performed in triplicate, and data are shown as mean ± standard deviation.

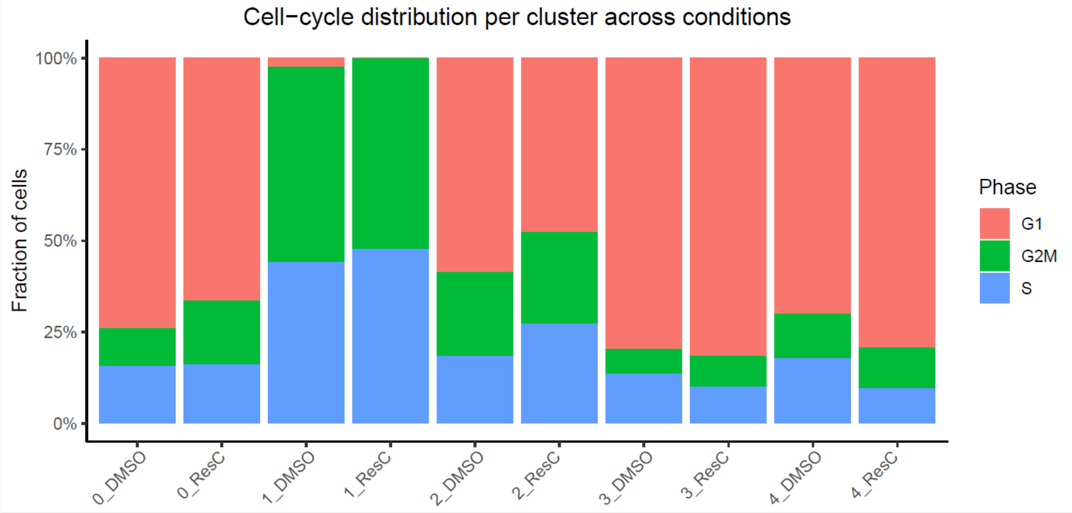


**Supplementary Fig 3. Cell-cycle composition of BI-2865–resistant clusters compared with DMSO controls in PDO-06.** Stacked bar charts illustrate the fractional distribution of cells across G1, S, and G2/M phases under each experimental condition. For individual clusters, cell-cycle profiles of BI-2865–resistant (ResC) populations are displayed alongside their matched DMSO-treated controls. Distinct, cluster-dependent remodeling of cell-cycle states is observed in resistant cells, characterized by differential representation of S-phase and G2/M compartments relative to control populations.

a b

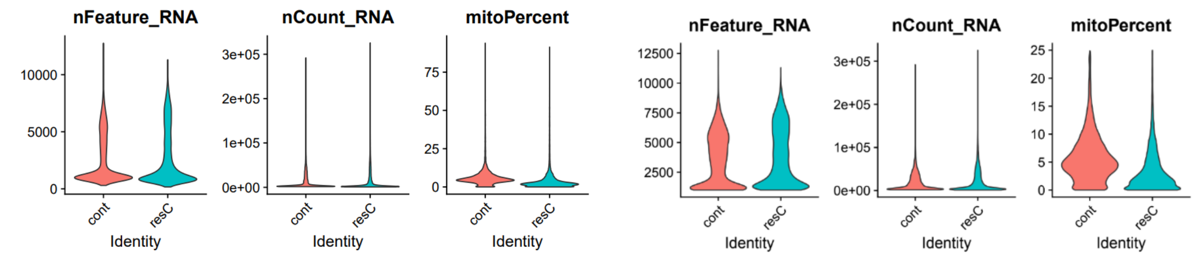


**Supplementary Fig 4. Quality control metrics before and after filtering of single-cell RNA sequencing data.**

Violin plots depict the distributions of the number of detected genes per cell (nFeature_RNA), total UMI counts per cell (nCount_RNA), and the percentage of mitochondrial transcripts (mitoPercent) across control (cont) and BI-2865–resistant (resC) populations. (a) Distributions prior to quality filtering. (b) Distributions after application of quality control thresholds, demonstrating removal of low-quality cells and normalization of mitochondrial content while preserving comparable transcriptional complexity between control and resistant cell.
